# Engineering a plant polyketide synthase for the biosynthesis of methylated flavonoids

**DOI:** 10.1101/2022.10.02.510496

**Authors:** Bo Peng, Lili Zhang, Siqi He, Rick Oerlemans, Wim J. Quax, Matthew R. Groves, Kristina Haslinger

## Abstract

Homoeriodictyol and hesperetin are naturally occurring O-methylated flavonoids with many health-promoting properties. They are produced in plants in low abundance and as complex mixtures of similar compounds that are difficult to separate. Synthetic biology offers the opportunity to produce various flavonoids in a targeted, bottom-up approach in engineered microbes with high product titers. However, the production of O-methylated flavonoids is currently still highly inefficient.

In this study, we investigated and engineered a combination of enzymes that had previously been shown to support homoeriodictyol and hesperetin production in *Escherichia coli* from fed O-methylated cinnamic acids. We determined the crystal structures of the enzyme catalyzing the first committed step of the pathway, chalcone synthase from *Hordeum vulgare*, in three ligand-bound states. Based on these structures and a multiple sequence alignment with other chalcone synthases, we constructed mutant variants and assessed their performance in *E. coli* towards producing methylated flavonoids. With our best mutant variant, HvCHS (Q232P, D234V), we were able to produce homoeriodictyol and hesperetin at 2 times and 10 times higher titers than previously reported. Our findings will facilitate the further engineering of this enzyme towards higher production of methylated flavonoids.

## 1. Introduction

Flavonoids are natural polyphenolic compounds ubiquitously found in various flowers, fruits, and vegetables^1,2^. In nature, flavonoids are important for plant growth and reproduction, for attracting pollinators, and for protecting against biotic and abiotic stresses, such as harmful ultraviolet radiation^3,4^. More than 15,000 compounds have been identified to date and many have been shown to have health-promoting effects^5^, such as anti-cancer, anti-inflammatory, anti-mutagenic, and anti-oxidative properties^6^. These health-promoting effects make flavonoids an attractive ingredient for nutraceutical, pharmaceutical, and cosmetic applications^1^.

Homoeriodictyol and hesperetin are important, naturally occurring O-methylated flavonoids, which exhibit higher biological activities and better pharmacological properties, including metabolic stability, membrane transport capability, and oral bioavailability, compared to unmethylated flavonoids^7^. Homoeriodictyol, which is O-methylated in the 3’-position, was previously isolated from *Viscum articulatum* Burm and has been shown to have anti-cancer effects in MCF-7, HeLa, and HT-29 cells^8^. Furthermore, it was shown to protect endothelial cells from oxidative stress and mitochondrial dysfunction^9^. Hesperetin, which is O-methylated in the 4’-position, has been reported to ameliorate Alzheimer’s disease^10^, and inhibit the migration of breast cancer^11^. More importantly, hesperetin displays strong inhibitory activity towards several viruses, such as influenza A 14 virus, parainfluenza virus type-3, and SARS-CoV^12,13^.

The current industrial production of flavonoids mainly relies on extraction from plants. This strategy has the limitations that the flavonoid content and yield is variable between seasons, and that flavonoids exist in complex mixtures that need to be separated with large volumes of organic solvents. The chemical synthesis of flavonoids requires harsh reaction conditions and toxic substrates, which are not eco-friendly^14^.

Another promising route for generating flavonoids is the biosynthesis of flavonoids in microbial cell factories. The rapid development of synthetic biology has enabled the heterologous expression of the flavonoid biosynthesis pathway in microorganisms such as *E. coli* and *Saccharomyces cerevisiae*^15–18^. In the first step, *p*-coumaric acid is activated by 4-coumarate-coenzyme A (CoA) ligase (4CL) to give coumaroyl-CoA (Fig. 1). Then, chalcone synthase (CHS) condenses one coumaroyl-CoA and three malonyl-CoA to form naringenin chalcone, which is subsequently converted into the (2S)-naringenin by chalcone isomerase (CHI). In plants, naringenin can be converted into other flavonoids by tailoring enzymes and is thus a key intermediate in the pathway. CHS is the first committed enzyme in flavonoid synthesis and belongs to the superfamily of type three polyketide synthases^19^. It has been reported that altering certain residues of such plant type three polyketide synthases improves their enzymatic activity and expands their substrate scope towards non-natural phenolic starter units^20,21^.

**Figure 1.**
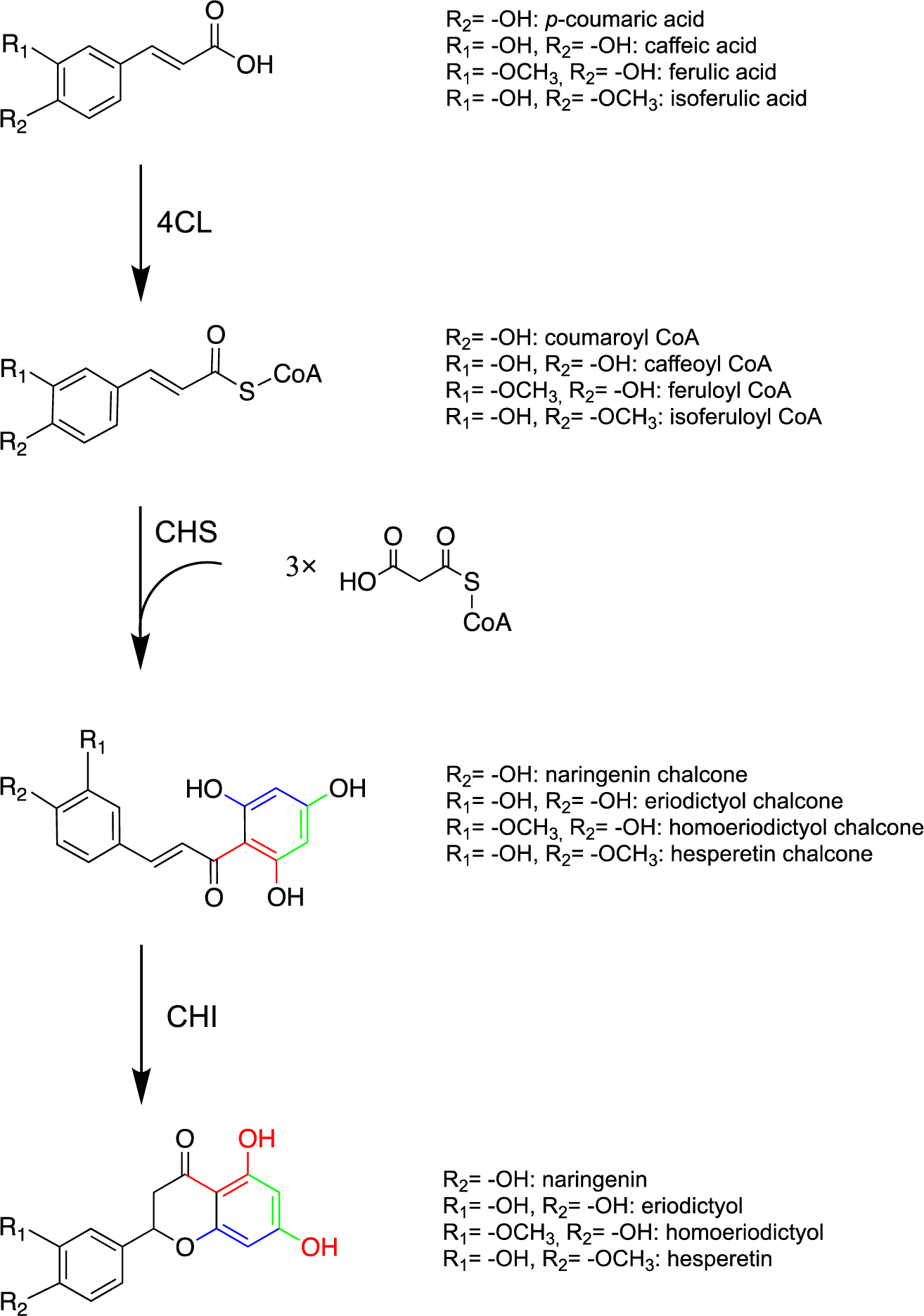
Biosynthesis pathway of the four flavonoids discussed in the study. The natural pathway product in plants is naringenin that is converted into the other flavonoids by tailoring enzymes. In this study, eriodictyol, homoeriodictyol and hesperetin are directly produced by feeding the respective precursors. (4CL 4-coumarate CoA ligase, CHS chalcone synthase, CHI chalcone isomerase).

Recently also other flavonoids were directly produced with this simple pathway in microbial cell factories by precursor feeding and pathway engineering. For instance, Dunstan *et al.* established high titers for naringenin (484 mg/L from *p*-coumaric acid), eriodictyol (55mg/L from caffeic acid), and homoeriodictyol (17mg/L from glycerol) with a semi-automated metabolic engineering strategy in *Escherichia coli*^16^. Cui *et al*. obtained low final titers of hesperetin (0.4mg/L) from isoferulic acid by constructing a recombinant *E. coli* strain that expresses 4CL from *Oryza sativa* (Os4CL) and CHS from *Hordeum vulgare* (HvCHS)^22^. Since efficient uptake of ferulic acid by *E. coli* has been demonstrated in other studies^23,24^, it is most likely that the low hesperetin titer in this study is related to overall poor enzymatic activity of this enzyme combination, or the fact that ferulic acid and isoferulic acid are poor substrates for these enzymes.

In this study, we set out to explore this enzyme combination further. We determined the crystal structure of HvCHS in complex with CoA, CoA and naringenin and CoA, and eriodictyol. Based on these structures, we designed mutant variants to increase the titers of the O-methylated flavonoids homoeriodictyol, and hesperetin produced in *E. coli* from fed ferulic acid and iso-ferulic acid.

## 2. Materials and methods

### 2.1 Bacterial strains, primers, and plasmids

All the bacterial strains and plasmids used in this study are listed in table 1 and table 2. Primers are listed in table S1. *E. coli* DH5a (New England Biolabs) was used for routine cloning and pathway propagation, *E. coli* MG1655 K-12 (DE3) was used for protein expression and fermentation, and *E. coli* BL21 (DE3) was used for protein purification^25^. The genes for PhCHS, Chalcone synthase from *Petunia hybrida*; MsCHI, Chalcone isomerase from *Medicago sativa*; Pc4CL, 4-coumarate--CoA ligase from *Petroselinum crispum*, HvCHS, Chalcone synthase from *Hordeum vulgare*; Os4CL, 4-coumarate--CoA ligase from *Oryza sativa* were codon optimized and synthesized from Integrate DNA Technology with restriction site introduced (Integrated DNA Technologies, Coralville, Iowa, USA). The genes for PhCHS and HvCHS were cloned between SacI and HindIII of pETDuet-1 and those for Pc4CL and Os4CL were cloned between NdeI and XhoI of pETDuet-1 yielding pETDuet-HvCHS-Os4CL and pETDuet-PhCHS-Pc4CL. The gene for MsCHI was cloned between NdeI and XhoI of pCDFDuet-1 and pETDuet-HvCHS yielding pCDFDuet-MsCHI and pETDuet-HvCHS-MsCHI. The gene for Os4CL was cloned between NdeI and XhoI of pCDFDuet-1 yielding pCDFDuet-Os4CL. The gene for HvCHS was cloned between ScaI and HindIII of pET28a yielding pET28a-HvCHS. The gene for Os4CL was cloned between NdeI and XhoI of pET28a yielding pET28a-Os4CL.

**Table 1.** List of plasmids and bacterial strains used in methylated flavonoids production strains.

| Name |  |  |  |  |  |
| --- | --- | --- | --- | --- | --- |
| Strains | Relevant Genotype |  |  | Reference |  |
| *E. coli* BL21(DE3) | F - ompT hsdSB(rB -mB - ) gal dem (λIts857 indI Sam7 nin5 lavUV5-T7gene1) | | | Invitrogen | |
| *E. coli* DH5a | supE44 $\Delta$(lacZYA-argF)U196 (Φ80ΔlacZM15) hsdR17 recA1 endA1 gyrA96 thi-1 relA1 | | | Invitrogen | |
| *E. coli* K-12 MG1655 (DE3) | K-12 F<sup>-</sup> $\lambda^-$ *ilvG<sup>-</sup> rfb-50 rph-1* | | | Nielsen *et al.*<sup>25</sup> | |

| Identifier | Plasmid name | Backbone | Enzyme encoded |  | Source |
| --- | --- | --- | --- | --- | --- |
|  |  |  | MCS1 | MCS2 |  |
| C1 | pETDuet::PhCHS_Pc4CL | pETduet-1 | PhCHS | Pc4CL | This study |
| C2 | pCDFDuet::_MsCHI | pCDFduet-1 | MsCHI | / | This study |
| C3 | pETDuet::HvCHS_Os4CL | pETduet-1 | HvCHS | Os4CL | This study |
| C4 | pETDuet::PhCHS_Pc4CL(del V342) | pETduet-1 | PhCHS | Pc4CL (del V342) | This study |
| C5 | pETDuet::HvCHS (A228S, D231I, Q232P, L233G, D234V)_Os4CL | pETduet-1 | HvCHS (A228S, D231I, Q232P, L233G, D234V) | Os4CL | This study |
| C6 | pETDuet::PhCHS_Pc4CL(Q214A) | pETduet-1 | PhCHS | Pc4CL (Q214A) |  |
| C7 | pETDuet::HvCHS_Os4CL (del V340) | pETduet-1 | HvCHS | Os4CL (del V340) | This study |
| C8 | pETDuet::HvCHS_Os4CL(Q212A) | pETduet-1 | HvCHS | Os4CL (Q212A) | This study |
| C9 | pETDuet::HvCHS_Os4CL(S242A) | pETduet-1 | HvCHS | Os4CL (S242A) | This study |
| C10 | pETDuet::HvCHS(A199T)_Os4CL | pETduet-1 | HvCHS (A199T) | Os4CL | This study |
| C11 | pETDuet::HvCHS(I267F)_Os4CL | pETduet-1 | HvCHS (I267F) | Os4CL | This study |
| C12 | pETDuet::PhCHS(T197A)_Pc4CL | pETduet-1 | PhCHS (T197A) | Pc4CL | This study |
| C13 | pETDuet::HvCHS(A228S)_Os4CL | pETduet-1 | HvCHS (A228S) | Os4CL | This study |
| C14 | pETDuet::HvCHS(D231I)_Os4CL | pETduet-1 | HvCHS (D231I) | Os4CL | This study |
| C15 | pETDuet::HvCHS(Q232P)_Os4CL | pETduet-1 | HvCHS (Q232P) | Os4CL | This study |
| C16 | pETDuet::HvCHS(L233G)_Os4CL | pETduet-1 | HvCHS (L233G) | Os4CL | This study |
| C17 | pETDuet::HvCHS(D234V)_Os4CL | pETduet-1 | HvCHS (D234V) | Os4CL | This study |
| C18 | pETDuet::HvCHS(Q232P, D234V)_Os4CL | pETduet-1 | HvCHS (Q232P, D234V) | Os4CL | This study |
| C19 | pETDuet::HvCHS(Q232P, D234V)_MsCHI | pETduet-1 | HvCHS (Q232P, D234V) | MsCHI | This study |
| C20 | pCDFDuet-1::Os4CL | pCDFDuet-1 | Os4CL | / | This study |
| C21 | pET28a::HvCHS | pET28a | HvCHS | / | This study |
| C22 | pET28a::HvCHS(Q232P, D234V) | pET28a | HvCHS (Q232P, D234V) | / | This study |
| C23 | pET28s::Os4CL | pET28a | Os4CL | / | This study |

\* MCS1 and 2, multiple cloning site (each with its own T7 promoter and terminator); PhCHS, CHS from *Petunia hybrida*; MsCHI, CHI from *Medicago sativa*; Pc4CL, 4CL from *Petroselinum crispum*; HvCHS, CHS from *Hordeum vulgare*; Os4CL, 4CL from *Oryza sativa*; codon optimized for expression in *E. coli*.

**Table 2.** List of *E. coli* MG1655 (DE3) strains used in fermentation experiments.

| Identifier | Plasmid name | Enzymes expressed | Source |
| --- | --- | --- | --- |
| S1 | C1, C2 | PhCHS, Pc4CL, and MsCHI | This study |
| S2 | C2, C3 | HvCHS, Os4CL, and MsCHI | This study |
| S3 | C2, C4 | PhCHS, Pc4CL (del V342), and MsCHI | This study |
| S4 | C2, C5 | HvCHS (A228S, D231I, Q232P, L233G, D234V), Os4CL, and MsCHI | This study |
| S5 | C2, C6 | PhCHS, Pc4CL (Q214A), and MsCHI | This study |
| S6 | C2, C7 | HvCHS, Os4CL (del V340), and MsCHI | This study |
| S7 | C2, C8 | HvCHS, Os4CL (Q212A), and MsCHI | This study |
| S8 | C2, C9 | HvCHS, Os4CL (S242A), and MsCHI | This study |
| S9 | C2, C10 | HvCHS (A199T), Os4CL, and MsCHI | This study |
| S10 | C2, C11 | HvCHS (I267F), Os4CL, and MsCHI | This study |
| S11 | C2, C12 | PhCHS (T197A), Pc4CL, and MsCHI | This study |
| S12 | C2, C13 | HvCHS (A228S), Os4CL, and MsCHI | This study |
| S13 | C2, C14 | HvCHS (D231I), Os4CL, and MsCHI | This study |
| S14 | C2, C15 | HvCHS (Q232P), Os4CL, and MsCHI | This study |
| S15 | C2, C16 | HvCHS (L233G), Os4CL, and MsCHI | This study |
| S16 | C2, C17 | HvCHS (D234V), Os4CL, and MsCHI | This study |
| S17 | C2, C19 | HvCHS (Q232P, D234V), Os4CL, and MsCHI | This study |
| S18 | C19, C20 | HvCHS (Q232P, D234V), MsCHI, and Os4CL | This study |

### 2.2 Synthesis of DNA template and protein mutagenesis

Mutagenesis libraries were generated using a modified QuikChange method^26^. The templates were digested with fast digest *Dpn*I overnight in 37°C incubators, then a PCR Clean-up kit was used to purify the QuikChange product. Subsequently, 1 µL of the purified DNA was transformed into *E. coli* DH5α. The plasmids were isolated with the QIAprep Spin Miniprep kit and were sequence verified.

### 2.3 Transformation of *E. coli* for methylated flavonoid production

To generate the flavonoid producing *E. coli* strains s1-s18, the respective plasmid combinations were co-transformed into *E. coli* MG1655 (DE3) electrocompetent cells. The protocol for preparation of electrocompetent cell was from Green *et al*^27^. After transformation, the cells were grown on selective LB agar (ampicillin and spectinomycin at 100 µg/mL) at 30°C overnight. The next day, colonies were inoculated in 3 mL selective LB liquid medium in round-bottom polystyrene tubes for cultivation at 30°C, 200 rpm overnight. The following day, glycerol stocks were prepared from these cultures (50% (v/v) glycerol) and stored at −80°C until further usage.

### 2.4 Multiple sequence alignment

Protein sequences were aligned with Clustal Omega (1.2.4) with default settings^28^. The alignment was visualized with AliView^29^. Active site residues of CHS were identified based on Protein Data Bank (PDB) entry 1CGK and those of 4CL based on 5BSW.

### 2.5 Fermentation conditions

For small-scale fermentation, the flavonoid pathway expressing strains were streaked in duplicates from glycerol stocks into LB agar plates with ampicillin and spectinomycin (100 µg/mL each). After incubation overnight at 30°C, single colonies were inoculated into a starter culture of 3 mL of selective LB medium in round-bottom polystyrene tubes and grown with shaking at 200 rpm at 30°C overnight. The next day, the cultures were diluted 1:100 into 3 mL seed cultures of modified MOPS medium^18^ for 24 h at 30°C. These were used to inoculate working cultures of 1 mL at an optical density at λ=600 nm (OD_600_) of 0.05 in a 48 flower-shaped well plate (m2p-lab, Germany) and incubated at 30°C, 900rpm in an Eppendorf Thermomixer. IPTG (final concentration, 1 mM), precursors (final concentration, 1 mM), and cerulenin (final concentration, 20 μg/mL) were added into the cell culture after 6 h and the incubation was continued for 32 h.

For large-scale fermentation, two single colonies of the respective strains were inoculated into starter cultures are described above. The next day, 1 mL of the starter culture was inoculated into 50 mL of fresh LB medium. IPTG (final concentration 1 mM) was added to the culture broths when the OD_600_ reached 0.4-0.6 for protein expression and the cultures were successively incubated at 30 °C for an additional 3 h. The cells were collected by centrifugation for 30 min at 3,700 rpm (Eppendorf 5920R, Germany) and 4 °C and then resuspended in 25 mL of fresh modified MOPS medium which included 1 mM precursor, 1 mM IPTG, and 20 μg/mL cerulenin for further 32 h fermentation^22^.

Samples were taken for the quantification of OD_600_ and extracellular metabolites after 32h fermentation^18^. Metabolites were prepared for HPLC and HPLC-MS analysis as described in Dunstan *et al.*^16^. Samples were stored in −20 °C until analysis. OD_600_ was measured by an absorbance plate reader (BMG Labtech, Germany) at 600 nm.

### 2.6 Analysis and quantification of target compounds

The authentic standards for *p*-coumaric acid, caffeic acid, ferulic acid, isoferulic acid, naringenin, eriodictyol, homoeriodictyol, hesperetin were purchased from Sigma Aldrich (USA). The compounds used in *in vitro* experiments (coenzyme A hydrate, malonyl coenzyme A tetralithium salt, and adenosine 5′-triphosphate (ATP) disodium salt hydrate) were purchased from Sigma Aldrich (USA). Fatty acid synthase (FAS) inhibitor cerulenin was purchased from Enzo life sciences (USA).

Samples from fermentation and *in vitro* turnover were analyzed by high performance liquid chromatography (HPLC), with a Shimadzu LC-10AT system equipped with a SPD-20A photodiode array detector (PDA). The samples were analyzed by 10μL injections and separation over an Agilent Eclipse XDB-C18 (5 μm, 4.6×150 mm) column with a concentration gradient (solution A: water + 0.1% trifluoroacetic acid (TFA), solution B: acetonitrile + 0.1% TFA) at a flow rate of 1 mL/min. The following gradient was used: 15% solution B for 3 min, 15-90% solution B over 6 min; 90% solution B for 2 min; 90-15% solution B over 3 min, 15% solution B for 4 min. *P*-coumaric acid, caffeic acid, ferulic acid, isoferulic acid, naringenin, eriodictyol, homoeriodictyol, and hesperetin were identified by comparison to authentic standards. The peak areas were integrated and converted to concentrations based on calibration curves with the authentic standards (Fig. S1).

The identity of hesperetin was furthermore confirmed by HPLC coupled mass spectrometry (HPLC-MS) with a Waters Acquity Arc UHPLC-MS equipped with a 2998 PDA, and a QDa single quadrupole mass detector (Fig. S2). The samples were separated over an XBridge BEH C18 3.5 µm column with a concentration gradient (solution A: water + 0.1% formic acid, and solution B: acetonitrile + 0.1% formic acid) at a flow rate of 0.25 mL/min (1 μL injections). The following gradient was used: 5% solution B for 2 min, 5-90% solution B over 3 min; 90% solution B for 2 min; 5% solution B for 3 min.

### 2.7 Protein expression, purification, and crystallization

#### Expression

*E. coli* BL21(DE3) harboring the plasmids pET28a::HvCHS, pET28a::HvCHS(Q232P, D234V) or pET28a::Os4CL were inoculated in 3 mL LB (containing 100μg ml^−1^ kanamycin) for overnight cultivation at 37°C, 200 rpm. The next day, 1 ml overnight cell culture was used to inoculate 1 L fresh self-induction medium for further 20 h cultivation at 30°C, 200 rpm in 5 L glass Erlenmeyer flask. The self-induction medium composition (1 L) consisted of 20 g tryptone, 5 g yeast, 5 g sodium chloride, 4.45 g disodium hydrogen phosphate dihydrate, 3.4 g potassium dihydrogen phosphate, 6 g glycerol, 0.5 g glucose, 1.28 g lactose, and 100 μg ml^−1^ Kanamycin.

#### Purification

All steps were performed with cooled buffers and at 4°C. Cells were harvested by centrifugation at 3,700 rpm. The cell pellet was resuspended in 20 mL lysis buffer (50 mM Tris-HCl, 300 mM NaCl, pH 7.6). The cells were disrupted by sonication for 4 × 40 s (with a 5 min rest interval between each cycle) at a 60 W output. The unbroken cells and debris were removed by centrifugation (18,000 rpm for 1h). The supernatant was filtered through a syringe filter (pore diameter 0.45 μm) and incubated with 3 mL Ni^2+^-sepharose resin, which had previously been equilibrated with lysis buffer, in a small column at 4°C for 18 h with agitation. The unbound proteins were eluted from the column using gravity flow. The column was first washed with lysis buffer (15 ml) and then with buffer A (30 ml, 50 mM Tris-HCl, 300 mM NaCl, 30 mM imidazole pH 7.6). Retained proteins were eluted with buffer B (5 ml, 50 mM Tris-HCl, 300 mM NaCl, 500 mM imidazole pH 7.6). Fractions were analyzed by separation using sodium dodecyl sulfate polyacrylamide gel electrophoresis (4–12% polyacrylamide) and staining with InstantBlue Coomassie Protein Stain (Abcam, UK). Fractions containing chalcone synthase were combined and loaded onto a HiLoad 16/600 Superdex 200 pg column, which had previously been equilibrated with buffer C (180 ml, 10 mM HEPES, 50 mM NaCl buffer, 5% glycerol, and 2 mM DTT, pH 8.5). Elution was performed by running buffer C across the column at 1 ml min^−1^ for 1.2 column volumes. Fractions were collected and analyzed by sodium dodecyl sulfate polyacrylamide gel electrophoresis. The purified enzyme was concentrated with centrifugal devices with Omega membrane 30K (Pall, USA) and stored at –80° C until further use.

#### Crystallization

Freshly prepared HvCHS was used in crystallization. The protein was concentrated to 10 mg/ml in a buffer consisting of 10 mM HEPES pH7.5; 50 mM NaCl; 5% (v/v) glycerol; 2 mM DTT. Screening for crystallization conditions was executed manually using commercial sparse-matrix screening kits (JCSG plus; PACT premier; Morpheus and the PGA screen; Molecular Dimensions Ltd.), by sitting drop vapor diffusion at 4°C. After 1-2 days, small crystals were obtained in a drop containing 1 μl protein and 1 μl crystallization buffer (0.1 M MES/Imidazole pH6.5; 0.03 M MgCl_2_, 0.03 M CaCl_2_; 20% (v/v) glycerol, 10% (v/v) PEG4000). The crystallization buffer was optimized to a lower concentration of precipitants (16% (v/v) glycerol and 8% (v/v) PEG4000) to get fewer and larger crystals (Fig. S3).

Crystals were harvested 3-4 days before diffraction using a nylon loop. Before flash-cooling in liquid nitrogen, crystals were quickly dipped into a cryoprotectant, composed of the reservoir buffer with an increased concentration of glycerol (32% (v/v)).

Some crystals were soaked in cryoprotectant containing either an additional 1 mM naringenin or 5 mM eriodictyol for 24 h in order to obtain crystals of the enzyme-product complex. Naringenin and eriodictyol were initially prepared as a 1 M stock in 100% DMSO.

#### Data collection, Structure determination and refinement

All diffraction data of HvCHS crystals were collected on beamlines P11 and P13 (operated by EMBL Hamburg) of Petra III at DESY (Hamburg, Germany)^30,31^.

Integration, space group determination, and scaling were carried out with XDSAPP and Aimless in the CCP4 suite^32,33^. The structure of HvCHS complexed with CoA was determined by molecular replacement with the monomer of chalcone synthase from *Arabidopsis thaliana* (PDB: 6DXB) using the program Phaser^34^. Molecular replacement with the DIMPLE pipeline^35^ was performed to determine and initially refine the structures of HvCHS in complex with naringenin and eriodictyol, utilizing the structure of HvCHS without ligands and waters as a starting model.

All structures were iteratively refined using manual adjustment in Coot^36^ and Refmac5^37^.

#### PDB deposition

The structures of HvCHS in complex with CoA, CoA and naringenin, and CoA and eriodictyol were deposited in the PDB under accession codes 8B32, 8B35, and 8B3C, respectively.

#### Structure comparison and visualization

The HvCHS structures were compared to known CHS structures in the PDB and pairwise to each other with the Dali server^38^. Final structures were visualized with PyMOL (Schrödinger, LLC).

### 2.8 *In vitro* synthesis of feruloyl-CoA

Feruloyl-CoA was synthesized in 5 mL *in vitro* reaction with purified Os4CL enzyme following established protocols^39,40^. The reaction mixture (purified enzyme (40 μg/ml), ferulic acid (400 μM), Coenzyme A (800 μM), ATP (2.5 mM), and MgCl_2_ (5 mM) in potassium phosphate buffer (50 mM, pH 7.4)) was incubated at 30°C in the dark, with mixing at 200 rpm overnight. Reaction product was analyzed by HPLC and then aliquoted and stored at −20°C for further experiments.

### 2.9 *In vitro* enzymatic assay

The *in vitro* enzymatic assays with HvCHS were performed in 50 µL reaction mixtures composed of feruloyl-CoA (2.5, 5, 10, 25, 50, 100 μM), malonyl-CoA (300 µM), and HvCHS variant (50 nM) in 100 mM phosphate buffer (pH 7.4). The individual reactions were started by adding the enzyme, incubated at 37°C without mixing, and quenched at different time points (2.5, 5, and 7.5 min) by adding an equal volume of methanol with 0.1% formic acid to quench the reaction. The samples were centrifuged at 15,000 rpm for 10 min, and the supernatant was used for HPLC analysis. The integrated peak areas of the product were converted into concentrations based on a calibration curve with the authentic standard, and plotted against time (Fig. S4). Apparent initial velocities were determined by linear regression over the early time points before 10% substrate conversion was reached. The apparent initial velocities of the biological triplicates were then plotted against the substrate concentrations and the resulting curves were fitted with the Michaelis-Menten equation in GraphPad prism. The full statistical analysis is provided in Table S2.

## 3. Results

Based on the results of previous studies on flavonoid production in *E. coli*, we chose two combinations of CHS and 4CL as a starting point for this study: PhCHS from *Petunia hybrida* and Pc4CL from *Petroselinum crispum*, previously shown to yield high titers of naringenin^15^, and HvCHS from *Hordeum vulgare* and Os4CL from *Oryza sativa*, previously shown to accept O-methylated cinnamic acids to form homoeriodictyol and hesperetin^22^. We cloned the genes into co-expression plasmids and transformed them into *E. coli* MG1655 (DE3) (Table 2). In an initial fermentation experiment at 1 mL scale, with fed ferulic acid, and isoferulic acid (1 mM final concentration), we observed that the combination of HvCHS and Os4CL (s2) yielded up to two times higher titers of homoeriodictyol than the other enzyme combination (s1) and that hesperetin was only produced by s2 (Fig. S5). Intrigued by this result, we wondered if mutagenesis of the 4CL enzymes to better accommodate the O-methylated substrates could further increase the titers of hesperetin and homoeriodictyol. Based on a previous mutagenesis study showing that the deletion of V341 in the *Nicotiana tabacum* 4CL allows for binding of double methylated sinapinic acid^41^, we deleted the corresponding residues in Pc4CL (V342) and Os4CL (V340). Furthermore, based on the crystal structure of the *Nicotiana tabacum* 4CL in complex with feruloyl-CoA (PDB: 5BSW)^41^, we hypothesized that mutating Q213 and S243 into alanine might provide more space in the substrate binding pocket to accommodate the O-methyl group. In Pc4CL, residue 243 is already an alanine. We screened the new 4CL mutants in the recombinant flavonoid pathway against the two methylated substrates and did not observe remarkable differences (Fig. S6) although the final homoeriodictyol titer with s6 (Os4CL delV340) is around 1.5 times higher than s2 for production (Fig. S6).

Thus, we proceeded with investigating the role of CHS in substrate selection by determining the crystal structures of HvCHS with the ligands CoA, naringenin, and eriodictyol by X-ray crystallography and by site-directed mutagenesis.

### 3.1 Crystal structure of chalcone synthase from *Hordeum vulgare*

The initial crystallization experiments with purified HvCHS yielded high-quality crystals that diffracted to 1.7 Å (Table 3). We solved the phase problem with molecular replacement using PDB: 6DXB ^42^ without the ligand. The refined structure (PDB: 8B32) shows high structural similarity with several CHS structures in the PDB based on a Dali search^43^ (Table S3). The highest Z-scores were determined for CHS1 from *Oryza sativa*, CHS from *Freesia hybrida*, (mutant variants of) CHS from *Medicago sativa* (MsCHS) and CHS1 from *Glycine max* (L.). These enzymes all share an amino acid sequence identity of more than 70% and adopt the typical CHS fold with the αβαβα pseudo-symmetric thiolase motif in the C-terminal domain and the α-helical motif in the N-terminal domain^44^ (Fig. 2A). In the refined structure, we observed additional electron density in the substrate binding cleft separating the N- and the C-terminal domain, which we interpreted as CoA. The putative CoA molecule occupies the area of the enzyme previously identified as the malonyl-CoA binding pocket^44^ and our interpretation of the electron density agrees well with the model of the ligand in the CoA-bound structure of MsCHS (PDB: 1BQ6) (Fig. 2B).

**Table 3.** Diffraction data collection, structure determination, and refinement statistics. Values in brackets refer to the highest resolution shell.

| Structure name |  | HvCHS+CoA | HvCHS+CoA, naringenin | HvCHS+CoA, eriodictyol |
| --- | --- | --- | --- | --- |
| PDB entry |  | 8B32 | 8B35 | 8B3C |
| Space group |  | P21 21 21 | P21 21 21 | P21 21 21 |
| Unit cell parameter | a,b,c<br>[Å] | a = 77.81 | a = 73.91 | a = 77.88 |
|  |  | b = 98.96 | b = 95.98 | b = 97.82 |
|  |  | c = 138.11 | c = 138.34 | c = 137.93 |
|  | α, β, γ<br>[°] | α = 90.00 | α = 90.00 | α = 90.00 |
|  |  | β = 90.00 | β = 90.00 | β = 90.00 |
|  |  | γ = 90.00 | γ = 90.00 | γ = 90.00 |
| Resolution, Å |  | 45.79-1.70 (1.73 - 1.70) | 47.99 - 2.00 (2.05- 2.00) | 48.91 - 2.00 (2.04 - 2.00) |
| Number of observations |  | 1556039 (8829) | 886223 (8504) | 921816 (8357) |
| Completness,% |  | 99.8 (97.7) | 99.9 (99.3) | 100 (98.7) |
| Multiplicity |  | 13.2 (13.7) | 13.2 (13.5) | 12.8 (12.5) |
| Rmerge |  | 0.07 (1.366) | 0.09 (0.476) | 0.08 (1.016) |
| Average I/σ,I |  | 18.3 (2.6) | 19.8 (6.4) | 17.0 (3.1) |
| CC(1/2) |  | 0.999 (0.885) | 0.999 (0.979) | 0.999 (0.947) |
| Refinement statistics |  |  |  |  |
| No. reflections, all/free |  | 117356/5895 | 67037/3335 | 71816/3554 |
| Rfactor |  | 0.177 | 0.159 | 0.198 |
| Rfree |  | 0.205 | 0.193 | 0.234 |
| No. protein atoms |  | 5859 | 5804 | 5838 |
| No. ligand atoms |  | 96 | 136 | 138 |
| No. water atoms |  | 401 | 367 | 164 |
| Average B-factors, Å² |  |  |  |  |
| Protein |  | 34.58 | 30.26 | 50.26 |
| Ligands |  | 55.21 | 56.41 | 83.47 |
| Water |  | 40.78 | 36.32 | 48.50 |
| rmsd from ideal values |  |  |  |  |
| Bond lengths, Å |  | 0.0129 | 0.0119 | 0.0094 |
| Bond angles, ° |  | 1.757 | 1.658 | 1.503 |
| Ramachandran plot |  |  |  |  |
| Favored, % |  | 96.97 | 96.31 | 97.11 |
| Allowed, % |  | 2.50 | 3.03 | 2.63 |

**Figure 2.**
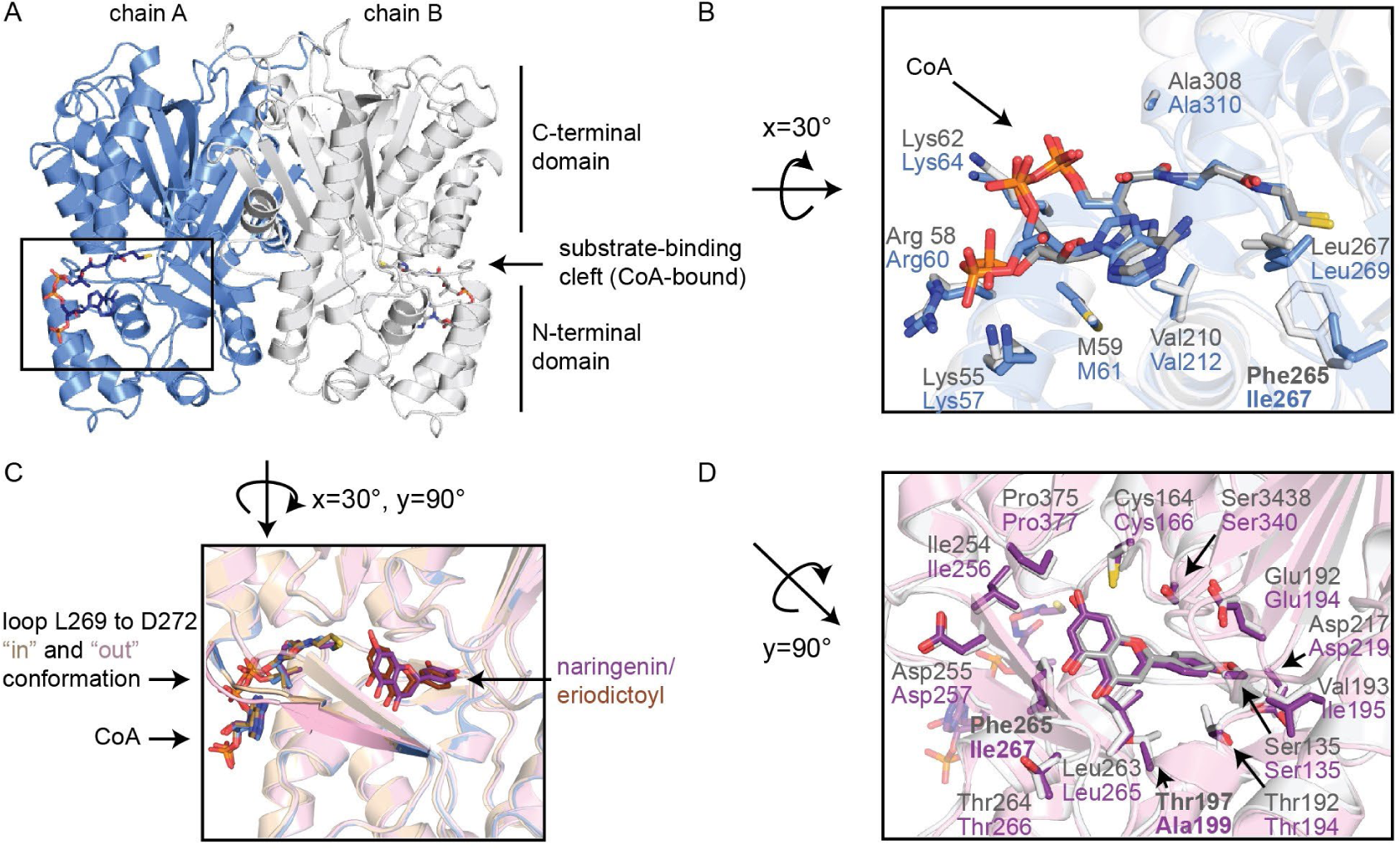
Crystal structures of HvCHS. A) Overview of HvCHS fold with the two monomers shown in grey and blue. B) CoA binding pocket of HvCHS in complex with CoA (blue) compared to the structure of MsCHS in complex with CoA (PDB: 1BQ6, grey). C) Overlay of all HvCHS structures (CoA-bound (blue), naringenin-bound (pink), eriodictoyl-bound (yellow). D) product binding pocket in HvCHS with naringenin bound (pink) compared to the structure of MsCHS in complex with naringenin (PDB: 1CGK, grey). Atom coloring in ligand representation: oxygen=red, nitrogen=blue, phosphorous=orange, carbon=color matches the cartoon color of the corresponding structure.

To gain further insight into the binding pocket of the cinnamic acid starter unit, we soaked the crystals of HvCHS with naringenin, eriodictyol, homoeriodictyol, and hesperetin. While the soaking with the O-methylated flavonoids impaired crystal diffraction, we were able to determine the structures of the naringenin-(PDB: 8B35) and eriodictyol-bound enzyme (PDB: 8B3C) at 2Å resolution. In both structures, the electron density for CoA and the products are well-defined and the position of the bound products agrees well with the MsCHS structure (PDB: 1CGK). All three HvCHS structures are highly congruent, even in residues lining the active site. The only difference can be seen in the β-sheet and loop from L269 to D272 (Fig 2C). Based on the electron density map, this loop appears to adopt two distinct conformations in all three crystal structures. However, while in the CoA-bound and the CoA/eriodictoyl-bound structures, the “in”-conformation has much higher occupancy and was therefore chosen for the final model, the “out”-conformation is dominant in the naringenin-bound structure. This agrees well with the conformation of this region in the naringenin-bound structure of MsCHS (1CGK, Fig 2D). This may be related to the slight difference in the angle of the A- and C-ring of naringenin and eriodictyol in the active site. The B-ring and all residues surrounding it, show identical conformations in both product-bound structures.

When comparing the structures and amino acid sequences of MsCHS and HvCHS, it becomes apparent that the residues in the substrate binding channel, and especially the cinnamoyl-binding pocket are highly conserved (Fig 2D, Fig. S7). The only differences are residue 197 in MsCHS, where a threonine is located compared to an alanine in HvCHS (residue 199), and residue 265, where a phenylalanine is located compared to an isoleucine in HvCHS (residue 267). Residue 197 is close to the C-ring of naringenin and may influence substrate binding. Mutation of residue 265 into valine was previously shown to decrease the efficiency of MsCHS two-fold but it did not alter the substrate scope of the enzyme^21^. Further away from the active site, we also noticed a small stretch of amino acids (228-234) with a markedly different sequence in HvCHS than in MsCHS and PhCHS. These residues are located on the surface of the enzyme in a loop that follows a β-strand, the opposite end of which forms part of the active site. Despite the dissimilarity in sequence, this loop adopts the same conformation in the MsCHS and HvCHS crystal structures (Fig. 28).

Based on our analysis of the sequence and structure of HvCHS, we decided to generate several mutant variants of HvCHS and PhCHS to probe the effect of these minor sequence differences on the enzyme performance for the production of methylated flavonoids *in vivo*.

### 3.2 Screening of CHS mutant variants for the production of methylated flavonoids

We constructed PhCHS T197A, HvCHS A199T, HvCHS I267F, and a HvCHS loop variant, where the sequence of the surface loop (228-234) is replaced by the sequence of the PhCHS loop. We transformed the resulting plasmids into *E. coli* to generate the flavonoid producing strains s4, s9, and s10 for the HvCHS variants and s11 for the PhCHS T197A variant. We performed small-scale fermentations (1 mL) of these strains feeding 1 mM ferulic acid or isoferulic acid as precursors, and sampled 32 h after induction of enzyme expression (Fig. 3A). The final optical densities (OD_600_) of all cultures were similar (ranging from 0.23 to 0.25) suggesting that the mutant variants do not affect the growth or viability of *E. coli*. Both point mutations in HvCHS (s9, s10) did not alter the final titers of homoeriodictyol compared to s2, yet the A199T mutation completely eliminated hesperetin formation in s9. The analogous point mutation in PhCHS (T197A, s11) for the first time enabled hesperetin production with this enzyme, while the homoeriodictyol titer from this strain is comparable to the strain expressing the wildtype protein (s1). Most interestingly, strain s4 expressing the HvCHS loop variant shows increased titers of both methylated flavonoids with final titers reaching 0.07 and 0.006 mM. These titers are about two-fold higher than the final titer achieved with HvCHS wildtype, and the hesperetin titer is comparable to the one achieved with the PhCHS T197A variant (s11). This suggests that the presence of alanine in this position in the active site enables the binding of isoferulic acid but it does not have a strong effect on ferulic acid binding. Furthermore, the residues in the surface loop appear to also influence the *in vivo* performance of the HvCHS enzyme, yet with an unknown mechanism.

**Figure 3.**
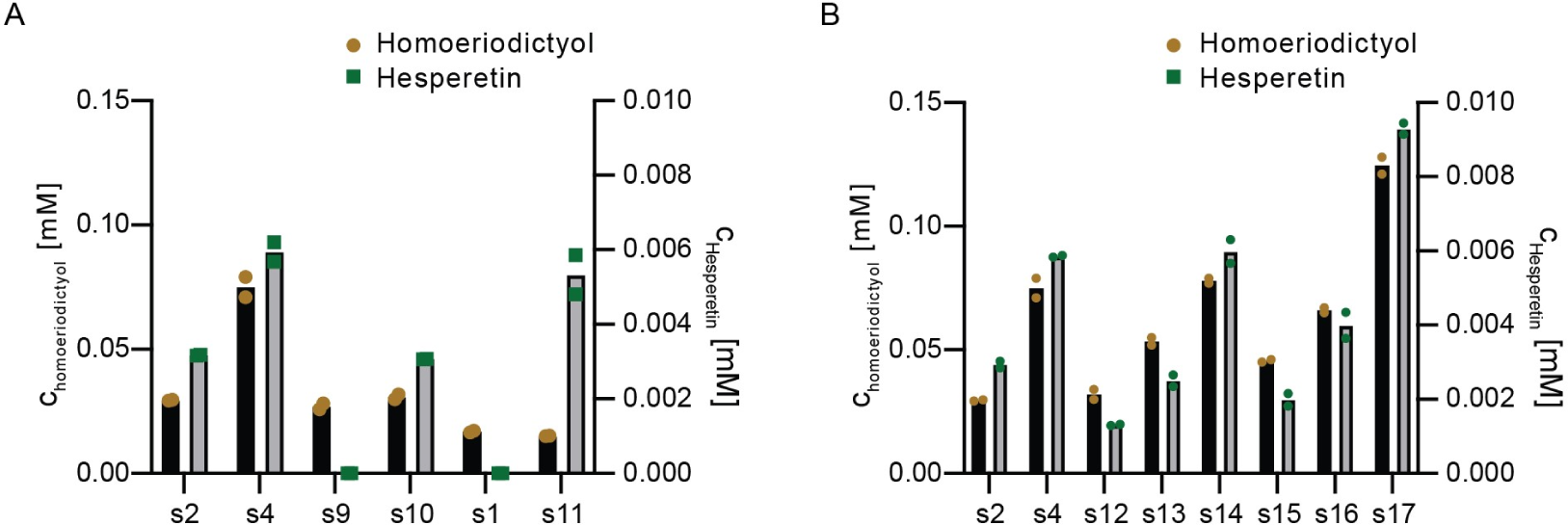
Small-scale fermentation of *E. coli* flavonoid producing strains expressing CHS variants. Titers of hesperetin (green) and homoeriodictyol (brown) were determined 32 h post induction of enzyme expression and addition of the precursors (1 mM ferulic acid or isoferulic acid). The experiment was performed in duplicate. A) Comparison of *E. coli* flavonoid producing strains expressing CHS variants: s1 PhCHS wildtype, s11 PhCHS T197A, s2 HvCHS wildtype, s4 loop variant, s9 HvCHS A199T, and s10 HvCHS I267F. B) Comparison of *E. coli* flavonoid producing strains expressing HvCHS variants: s2 wildtype, s4 loop variant, s12-s16 single point mutation variants, s17 double point mutation variant.

In order to further probe the importance of the individual residues of the surface loop for the enhanced *in vivo* performance of the HvCHS loop variant (A228S, D231I, Q232P, L233G, D234V), we next constructed single mutants for those five sites via site-directed mutagenesis. We transformed the resulting plasmids into *E. coli* (s12-s16) and compared the final titers of fermentations with these strains to the ones expressing HvCHS wildtype and the loop variant (s2 and s4, respectively). Compared to s4, only s14 and s16 yielded similar or higher product titers (Fig. 3B). S12 produced lower and s13 and s15 similar titers as s2 expressing wildtype HvCHS. Therefore, we combined the two best point mutations into a double mutant HvCHS (Q232P, D234V) and transformed the resulting plasmid into *E. coli* (s17). This strain yields higher titers of the methylated flavonoids than all other strains, with final homoeriodictyol titers 2-fold higher, and final hesperetin titers 3-fold higher than with strain s2 expressing wildtype HvCHS (Fig. 3B).

### 3.3 Further optimization of flavonoid production

To test if we could further increase the titers of methylated flavonoids with our HvCHS variant, we explored a different plasmid configuration. Santos *et al*. reported that it was optimal for naringenin production to clone CHS into multiple cloning site (MCS) 1 of pETDuet-1, MsCHI into MCS2 and 4CL into MCS1 of pCDFduet-1^15^. Thus, we constructed expression plasmids c19 and c20 with our best mutant variant according to this design and co-transformed them into *E. coli* (s18). We performed a fermentation at larger scale in shake flasks to see how the titers for methylated flavonoids from s18 compare to those of s2 and s17 (Fig. 4A). We found that the final titers of both products from s18 are comparable to those obtained with s2 (0.24 mM for homoeriodictyol and 0.011 mM for hesperetin). The final titer of homoeriodictyol and hesperetin are highest with s17, 0.33 mM and 0.016 mM, respectively. We next examined the production of two unmethylated flavonoids, naringenin and eriodictyol, from their respective cinnamic acid precursors with our best strain in shake flasks. After 32 h, we achieved titers of 41 mg/L and 45 mg/L, respectively. Overall, our best strain s17 produces similar or higher titers of the four flavonoids compared to the study of Cui *et al.*^22^.

**Figure 4.**
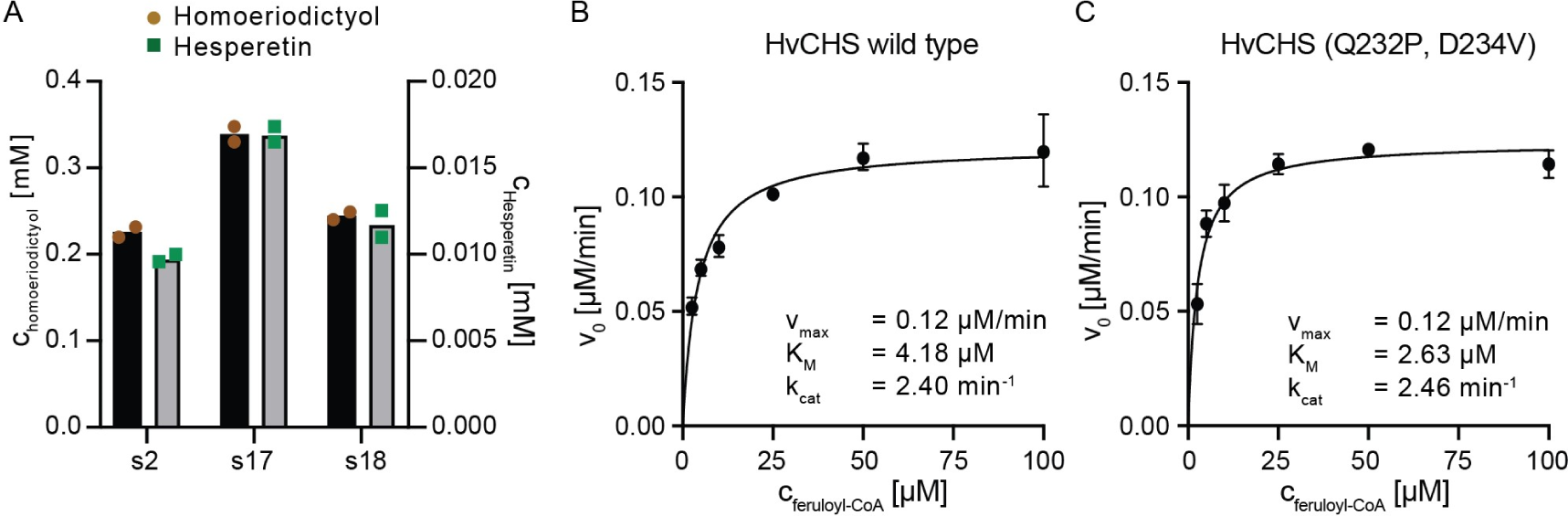
A) Comparison of HvCHS wildtype and double mutant for methylated flavonoid production in a large-scale fermentation. 1 mM ferulic acid and isoferulic acid were added as precursors. Samples were taken after 32 h fermentation and analyzed by HPLC. Each experiment was duplicated. HvCHS variants *in vitro* kinetic data. B-C) Kinetic characterization of HvCHS wildtype and HvCHS double mutant for homoeriodictyol formation. Apparent initial velocities were determined at 5, 10, 25, 50, and 100 µM substrate concentration (feruloyl-CoA). Data points represent mean +/− SD, n=3. The full statistical report for non-linear regression is shown in Table S2.

### 3.4 Enzymatic assay for HvCHS variants

Intrigued by the observation that the surface loop residues of HvCHS have an impact on the enzyme’s *in vivo* performance, we endeavored to further examine this phenomenon *in vitro*. We expressed and purified HvCHS wildtype and double mutant (Q232P, D234V) with protein yields of 48 and 93 mg protein/ L culture, respectively. This suggests that the double mutant has a higher expression level, is more soluble, or more stable during purification than the wildtype enzyme. Given that homoeriodictyol is the main product *in vivo*, we performed steady-state kinetic assays for wildtype and double mutant using feruloyl-CoA as a substrate at a fixed malonyl-CoA concentration. Plotting the apparent initial reaction velocities over the substrate concentration allowed us to fit the data with the Michaelis-Menten equation (Fig. 4B/C, Fig. S4, Table S2). The apparent kinetic parameters v_max_ and k_cat_ are virtually the same for both HvCHS variants, whereas there is a 1.6 fold difference in the K_M_. This indicates that the HvCHS (Q232P, D234V) variant exhibits a higher affinity towards feruloyl-CoA than the wildtype and is in good agreement with our results from homoeriodityol production *in vivo*. In summary, the enhanced biosynthesis of flavonoids (naringenin, eriodictyol, homoeriodictyol, and hesperetin) in *E. coli* expressing the HvCHS double mutant (Q232P, D234V) can likely be attributed to the combined effects of a higher protein expression level, and a higher affinity to methylated substrates.

## 4. Discussion and Conclusions

Hesperetin and homoeriodictyol are valuable natural O-methylated flavonoids which display bioactivity for treating human disease, such as neuroinflammation, memory impairment and carcinoid cancer^45,46^. Recently, Cui *et al*. obtained hesperetin and homoeriodictyol at a low titer in engineered *E. coli* from fed isoferulic acid and ferulic acid^22^. The enzymes used in that study had previously been shown to have high substrate promiscuity *in vitro*^47,48^ and were then used *in vivo* for the first time^22^. To become industrially relevant, the yield and productivity of such a pathway must be dramatically increased and enzymes must be highly selective for the O-methylated substrates to allow for the use of low-cost substrates. Thus, to facilitate further enzyme engineering efforts, we determined the crystal structures of one of the enzymes of this pathway, HvCHS, in complex with its products and analyzed a multiple sequence alignment with the most similar structures in the PDB. Since all CHS enzymes are highly conserved overall, especially in the substrate binding pockets, we did not see any obvious reasons why HvCHS would accept O-methylated precursors and other CHS enzymes would not. However, we identified three areas of interest for rational enzyme engineering – two amino acids in the substrate binding pocket that are markedly different from the consensus sequence (A199 and I267 in HvCHS) and a surface exposed loop that is connected to the active site through a β-strand and has more polar residues than in the other CHS sequences. Indeed, mutating the conserved Thr at position 197 in PhCHS enabled this enzyme to produce hesperetin for the first time. However, replacing I267 with the canonical Phe did not alter the performance of HvCHS. Surprisingly, mutating two of the polar or charged residues of the surface loop into hydrophobic amino acids increased the final titers of methylated flavonoids *in vivo*. We were able to show that this is likely attributable to a boost in protein abundance and an increase in affinity for the methylated cinnamic acid.

The final titers that we achieved for both methylated flavonoids in a larger scale experiment, exceed the previously reported titers by 2- and 10-fold for homoeriodictyol and hesperetin production. Compared to homoeriodictyol, hesperetin production is still very low, mainly due to the substrate preference of HvCHS. In the future, this substrate preference can be further shifted by directed evolution or rational engineering based on our crystal structure. In particular, in light of our surprising findings about the surface loop distant to the active site, it is worth exploring other positions in the multiple sequence alignment that are markedly different in HvCHS compared to other CHS enzymes. Alternatively, as more than 1000 putative plant CHSs have now been predicted via computational tools, it is also interesting to investigate the substrate scope and catalytic activity of these new enzymes to possibly replace the workhorse CHS enzymes with ones that are better suited for large-scale production of hesperetin and other flavonoids. Furthermore, an alternative strategy could involve employing host strains that possess an ample supply of malonyl-CoA for enhanced flavonoid synthesis.

## Abbreviations and Nomenclature

4CL: 4-coumarate: CoA ligase
CHI: chalcone isomerase
CHS: chalcone synthase
CoA: Co-enzyme A
HPLC: High Performance Liquid Chromatography
MS: mass spectrometry
TFA: trifluoroacetic acid

## Supporting Information

Gene and primer sequences, results of Dali analysis, statistical analysis of steady-state kinetics, calibration curves for product quantification by HPLC, HPLC-MS identification of fermentation products, micrograph of protein crystals, time progress curves for steady-state kinetics, product titers of fermentations with strains s1, s2, s3, s5, s6, and s7 expressing 4CL variants, sequence comparison of CHS enzymes, illustration of HvCHS surface loop (PDF).

## Acknowledgement

The authors acknowledge DESY (Hamburg, Germany) and EMBL Hamburg for the provision of the PETRA III beamline (P11 and P13). KH, BP, SH and WJQ thank Serj Koshian for designing the initial set of mutant enzymes during his summer internship, Dr. Robbert Hans Cool for providing training in protein purification, and Nika Sokolova for help with troubleshooting the steady-state kinetics.

## Author contributions

KH and BP conceived the study with contributions from LZ, SH, RO, WJQ and MG; BP and SH cloned plasmids and performed fermentations; BP and KH analyzed biochemical data; LZ and RO performed crystallization and diffraction experiments; LZ, RO and MG determined and refined the crystal structures; KH and BP wrote the manuscript with contributions from LZ, SH, RO, WJQ and MG; all authors have read and approved the final version of the manuscript.

## For table of contents only

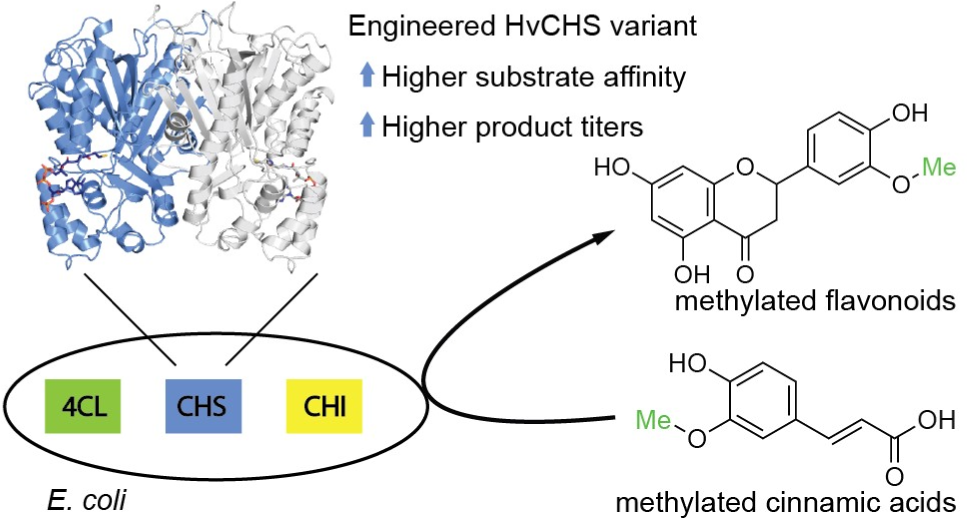

