## Supplementary Figures and Tables for "Engineering a plant polyketide synthase for the biosynthesis of methylated flavonoids"

### Supporting Materials

#### *Sequences of synthetic genes*

Pc4CL (*Petroselinum crispum*,  GenBank accession number KX671122.1):

ATGGGTGACTGCGTTGCCCCGAAAGAGGATCTGATCTTCCGCAGCAAACTGCCGGACATTTACATTCCAAAGCATCTGCCGCTGCATACC
TATTGTTTTGAGAACATCAGCAAGGTTGGCGACAAGAGCTGTCTGATCAACGGCGCAACCGGCGAAACCTTTACCTACAGCCAGGTTGA
GCTGCTGTCCCGTAAAGTTGCCAGCGGCCTGAACAAGCTGGGCATTCAACAAGGTGATACCATTATGCTGCTGCTGCCGAACTCCCCGG
AGTACTTTTTCGCTTTCCTGGGTGCGAGCTATCGCGGTGCAATCAGCACTATGGCGAACCCATTCTTTACCAGCGCAGAAGTGATCAAGC
AACTGAAAGCGAGCCAAGCGAAGCTGATTATCACCCAGGCATGCTATGTTGACAAGGTTAAGGACTACGCAGCGGAGAAAAACATCCA
GATCATTTGTATTGACGATGCACCGCAGGATTGCCTGCACTTTAGCAAGCTGATGGAAGCGGATGAGAGCGAAATGCCGGAAGTGGTT
ATTAACAGCGATGATGTGGTGGCACTGCCGTACAGCTCTGGCACCACCGGCCTGCCGAAAGGCGTTATGCTGACCCACAAGGGTCTGGT
TACCAGCGTTGCACAACAGGTGGATGGTGATAACCCGAACCTGTATATGCACTCCGAGGATGTTATGATCTGCATCCTGCCACTGTTCCA
TATCTATAGCCTGAACGCTGTTCTGTGTTGTGGTCTGCGTGCGGGCGTTACCATTCTGATCATGCAAAAGTTCGACATTGTGCCGTTTCTG
GAGCTGATTCAGAAGTATAAGGTTACCATTGGTCCGTTTGTTCCGCCGATCGTGCTGGCCATCGCGAAAAGCCCGGTTGTTGACAAGTA
CGACCTGTCTAGCGTGCGCACCGTTATGAGCGGTGCAGCGCCGCTGGGTAAAGAGCTGGAGGACGCTGTTCGTGCGAAATTCCCGAAC
GCGAAGCTGGGTCAAGGCTATGGCATGACCGAAGCCGGTCCGGTTCTGGCGATGTGTCTGGCGTTCGCCAAAGAGCCGTATGAGATTA
AGTCTGGCGCATGCGGTACCGTTGTGCGTAACGCCGAGATGAAAATCGTTGACCCAGAAACCAACGCGTCTCTGCCGCGTAACCAGCGT
GGTGAGATTTGCATCCGTGGTGATCAGATTATGAAAGGTTACCTGAACGACCCGGAAAGCACCCGCACCACCATCGACGAAGAGGGTT
GGCTGCACACCGGTGACATTGGTTTCATCGACGATGACGATGAACTGTTCATTGTTGATCGTCTGAAAGAAATCATTAAGTACAAAGGTT
TTCAAGTTGCTCCGGCGGAGCTGGAAGCACTGCTGCTGACCCACCCGACCATCAGCGATGCCGCGGTGGTTCCGATGATTGACGAGAA
AGCGGGTGAAGTGCCAGTGGCGTTTGTTGTGCGTACCAACGGTTTTACCACCACCGAAGAAGAAATCAAACAATTTGTGAGCAAACAG
GTTGTGTTCTACAAACGTATCTTCCGCGTTTTCTTCGTTGACGCTATTCCGAAATCCCCGAGCGGCAAGATTCTGCGTAAGGATCTGCGC
GCTCGTATTGCGAGCGGCGACCTGCCGAAGTAA

Os4CL (*Oryza sativa*, GenBank accession number NP_001396278):

ATGGGGTCAGTTGCAGCCGAGGAAGTAGTCGTCTTCCGCTCTAAGCTGCCGGATATCGAAATCGATAACTCTATGACCCTGCAAGAATA
CTGCTTTGCACGTATGGCAGAGGTGGGAGCCCGCCCATGTCTTATCGACGGGCAAACTGGAGAGTCATACACTTATGCTGAAGTAGAAT
CGGCCTCGCGTCGCGCAGCCGCGGGACTTCGTCGCATGGGAGTGGGCAAGGGAGACGTAGTGATGTCACTTTTGCGCAATTGCCCAGA
GTTTGCCTTTTCTTTTCTGGGGGCTGCCCGCTTAGGAGCGGCTACTACGACTGCAAATCCATTTTATACACCCCATGAGGTACATCGCCAG
GCTGAGGCAGCGGGGGCACGCGTTATTGTGACGGAAGCCTGCGCAGTTGAAAAGGTGCGCGAGTTCGCAGCTGAACGTGGTGTGCCT
GTTGTGACTGTCGACGGTGCCTTTGATGGGTGCGTAGAATTTCGTGAAGTGCTTGCAGCGGAGGAGTTAGATGCTGACGCTGATGTCCA
CCCCGATGATGTAGTGGCGCTTCCTTATTCTTCGGGTACCACAGGCTTACCTAAGGGCGTCATGCTGACACATCGTTCCCTTATTACATCG
GTTGCGCAGCAAGTAGATGGGGAAAACCCTAATCTTTATTTCTCCAAAGATGACGTGATTTTATGCCTGCTGCCTCTTTTTCATATTTATT
CGTTAAACAGCGTTCTGCTGGCAGGGCTGCGCGCTGGTTCTACAATTGTCATCATGCGCAAATTTGATTTGGGGGCGCTTGTTGATCTGG
TCCGTAAGCATAACATTACGATTGCACCATTTGTACCACCCATCGTTGTAGAAATTGCTAAATCACCCCGCGTAACAGCTGAGGATCTGG
CCTCTATTCGTATGGTCATGTCAGGTGCAGCTCCTATGGGTAAGGATTTGCAGGACGCCTTTATGGCGAAAATCCCTAATGCAGTTTTAG
GTCAAGGGTATGGTATGACTGAGGCAGGCCCTGTTCTGGCAATGTGTCTGGCCTTCGCCAAAGAGCCGTTTAAGGTAAAGTCCGGAAG
CTGTGGTACCGTGGTTCGCAACGCTGAACTTAAGATCGTAGATCCCGACACGGGTACGTCGCTGGGTCGCAATCAATCTGGTGAGATCT
GCATCCGTGGTGAGCAGATTATGAAGGGCTACTTAAACGATCCAGAGGCTACAAAGAACACCATCGATGAGGATGGGTGGCTGCATAC
CGGAGACATTGGTTTCGTGGACGATGATGATGAGATTTTTATTGTCGACCGTCTTAAAGAAATTATCAAGTACAAGGGGTTCCAGGTAC
CACCCGCTGAATTGGAGGCTCTTCTTATCACTCACCCAGAGATCAAGGACGCGGCGGTAGTCAGCATGAAGGACGATTTGGCGGGAGA
GGTCCCTGTAGCCTTCATTGTCCGCACTGAGGGGAGCGAAATTACAGAGGACGAAATTAAAAAGTTCGTTGCCAAAGAGGTGGTGTTCT
ACAAGCGTATTAACAAAGTGTTCTTTACAGATTCCATCCCTAAAAACCCTTCCGGTAAGATCTTGCGCAAGGATCTTCGTGCTCGTCTGGC
GGCGGGTATTCCGGACGCTGTCGCTGCGGCAGCTGCAGATGCTCCTAAAAGTAGCTAA

HvCHS (*Hordeum vulgare*, GenBank accession number XP_044963194):

ATGGCAGCGGTGCGTTTGAAGGAGGTGCGCATGGCGCAGCGCGCCGAGGGTTTAGCTACAGTGCTGGCGATCGGAACGGCTGTACCT
GCAAATTGTGTTTACCAAGCGACATATCCTGACTACTACTTTCGTGTTACTAAGTCAGAGCATTTAGCCGATCTTAAAGAAAAGTTTCAGC
GCATGTGTGACAAGAGTATGATCCGCAAACGCCACATGCACCTTACGGAAGAGATTTTAATTAAGAACCCCAAAATTTGCGCCCACATG
GAGACTTCACTTGATGCCCGTCATGCCATTGCCTTGGTGGAAGTCCCGAAATTGGGCCAAGGGGCCGCCGAAAAAGCCATTAAAGAAT
GGGGCCAACCGCTTAGTAAGATCACGCACTTGGTCTTTTGCACAACATCAGGAGTTGATATGCCCGGGGCGGATTACCAGTTAACGAAG
CTCCTGGGATTGAGCCCTACTGTCAAACGCCTTATGATGTACCAACAAGGATGTTTTGGCGGAGCTACTGTATTACGCCTGGCCAAAGAT
ATTGCCGAGAACAATCGCGGGGCTCGCGTTTTAGTAGTTTGCTCGGAGATTACCGCAATGGCGTTCCGTGGCCCGTGCAAATCCCATTT
GGATTCGTTAGTAGGTCACGCATTATTCGGCGATGGAGCCGCTGCTGCAATCATCGGAGCCGATCCCGACCAATTAGACGAGCAGCCG
GTATTTCAATTGGTATCGGCGAGTCAGACAATCCTGCCAGAATCGGAGGGTGCGATCGATGGACACTTGACGGAGGCGGGCTTAACGA
TTCACCTGCTGAAAGATGTGCCGGGCTTGATCTCTGAAAACATCGAGCAGGCTCTTGAGGATGCGTTTGAGCCTTTGGGTATTCACAATT
GGAACTCTATTTTTTGGATCGCTCATCCGGGCGGGCCTGCAATTTTAGACCGCGTCGAAGATCGCGTTGGATTAGATAAAAAACGTATG
CGTGCTTCACGCGAGGTCCTTAGCGAATACGGCAACATGTCTTCAGCCTCTGTCTTATTCGTGCTTGACGTTATGCGCAAGTCGAGTGCA
AAGGATGGGTTGGCGACGACGGGCGAGGGGAAAGATTGGGGGGTGCTGTTCGGATTTGGCCCAGGACTGACCGTCGAAACCCTGGTA
TTACACTCTGTTCCTGTTCCCGTTCCCACTGCGGCTTCTGCTTAA

PhCHS (*Petunia hybrida*, GenBank accession number KP284563.1 ):

ATGGTGACCGTGGAAGAATACCGTAAGGCGCAACGTGCGGAAGGCCCGGCGACCGTGATGGCGATTGGCACCGCGACCCCGAGCAAC
TGCGTTGACCAGAGCACCTACCCGGATTTCTATTTTCGTATTACCAACAGCGAGCACAAAACCGACCTGAAGGAAAAATTCAAGCGTAT
GTGCGAGAAGAGCATGATTAAGAAACGTTACATGCACCTGACCGAGGAAATCCTGAAAGAGAACCCGAGCATGTGCGAATATATGGCG
CCGAGCCTGGACGCGCGTCAGGATATCGTGGTTGTGGAAGTGCCGAAACTGGGCAAAGAGGCGGCGCAGAAAGCGATTAAGGAATG
GGGTCAACCGAAAAGCAAGATCACCCACCTGGTTTTCTGCACCACCAGCGGCGTGGACATGCCGGGTTGCGATTACCAACTGACCAAAC
TGCTGGGCCTGCGTCCGAGCGTTAAGCGTCTGATGATGTATCAGCAAGGTTGCTTTGCGGGTGGCACCGTGCTGCGTCTGGCGAAAGA
TCTGGCGGAAAACAACAAGGGTGCGCGTGTTCTGGTTGTGTGCAGCGAGATTACCGCGGTGACCTTCCGTGGCCCGAACGACACCCAC
CTGGATAGCCTGGTTGGTCAGGCGCTGTTTGGTGATGGTGCGGGTGCGATCATTATCGGCAGCGATCCGATTCCGGGTGTTGAGCGTCC
GCTGTTCGAACTGGTGAGCGCGGCGCAAACCCTGCTGCCGGACAGCCATGGTGCGATTGATGGTCACCTGCGTGAAGTTGGCCTGACC
TTTCACCTGCTGAAAGACGTGCCGGGTCTGATTAGCAAAAACATCGAGAAGAGCCTGGAGGAAGCGTTCAAGCCGCTGGGCATTAGCG
ACTGGAACAGCCTGTTTTGGATTGCGCACCCGGGTGGCCCGGCGATTCTGGATCAAGTTGAAATCAAACTGGGCCTGAAGCCGGAGAA
ACTGAAGGCGACCCGTAACGTTCTGAGCAACTACGGTAACATGAGCAGCGCGTGCGTGCTGTTTATCCTGGATGAAATGCGTAAAGCG
AGCGCGAAAGAGGGTCTGGGTACCACCGGCGAGGGTCTGGAATGGGGTGTGCTGTTCGGCTTTGGTCCGGGCCTGACCGTGGAAACC
GTTGTTCTGCATAGCGTTGCGACCTAA

MsCHI (*Medicago sativa*, GenBank accession number P28012):

ATGGCGGCGAGCATTACCGCGATTACCGTGGAAAATCTGGAATATCCGGCGGTTGTGACCAGCCCGGTGACCGGCAAAAGCTACTTCCT
GGGTGGCGCGGGCGAGCGTGGCCTGACCATCGAAGGCAACTTCATTAAATTTACCGCGATCGGTGTGTACCTGGAGGACATTGCGGTT
GCGAGCCTGGCGGCGAAGTGGAAAGGCAAGAGCAGCGAGGAACTGCTGGAAACCCTGGACTTCTATCGTGATATCATTAGCGGTCCGT
TTGAAAAACTGATTCGTGGCAGCAAGATCCGTGAGCTGAGCGGTCCGGAATACAGCCGTAAAGTGATGGAGAACTGCGTTGCGCACCT
GAAGAGCGTGGGTACCTATGGCGATGCGGAGGCGGAAGCGATGCAGAAATTCGCGGAAGCGTTTAAACCGGTGAACTTCCCGCCGGG
TGCGAGCGTGTTCTACCGTCAAAGCCCGAACGGTATTCTGGGCCTGAGCTTCAGCCCGGACACCAGCATTCCGGAGAAAGAAGCGGCG
CTGATCGAAAACAAGGCGGTGAGCAGCGCGGTTCTGGAAACCATGATCGGTGAACACGCGGTTAGCCCGGATCTGAAGCGTTGCCTGG
CGGCGCGTCTGCCGGCGCTGCTGAATGAGGGTGCGTTCAAGATTGGTAACTAA

### Supporting Tables

**Table S1.** Primers used in this experiment

| Primers | Templates | Primer sequences (5′ to 3′, mutant site underlined) | Products |
|---|---|---|---|
| Pc4CL del V342 FP<br>Pc4CL RP | C1 | AAGCCGGTCCGCTGGCGATGTG<br>GTGGTGCGGGTGCTTTCCGG | C6 |
| Pc4CL Q214A FP<br>Pc4CL RP | C1 | CAGCGTTGCACAAGCAGTGGATGGTGAT<br>GTGGTGCGGGTGCTTTCCGG | C8 |
| HvCHS(A228S, D231I, Q232P, L233G, D234V) FP<br><br>HvCHS (A228S, D231I, Q232P, L233G, D234V) RP | C3 | [PHO]GGGTGTTGAGCAGCCGGTATTTCAATTGGTATCGGCGAGTC<br>[PHO]GGGATGGGATCGGATCCGATGATTGCAGCAGC | C7 |
| Os4CL del V340 FP<br>Os4CL del V340 RP | C3 | CTGAGGCAGGCCCTCTGGCAATGTGTCT<br>AGACACATTGCCAGAGGGCCTGCCTCAG | C9 |
| Os4CL Q212A FP<br>Os4CL Q212A RP | C3 | ATTACATCGGTTGCGCAGGCGGTAGATGGGGAAAACCC<br>GGGTTTTCCCCATCTACCGCCTGCGCAACCGATGTAAT | C10 |
| Os4CL S242A FP<br>Os4CL S242A RP | C3 | ATTTATTCGTTAAACGCGGTTCTGCTGGCAGGG<br>CCCTGCCAGCAGAACCGCGTTTAACGAATAAAT | C11 |
| HvCHS A199T FP<br>HvCHS A199T RP | C3 | TTACCGCAATGACCTTCCGTGGCCCGT<br>GCCACGGAACGCCATGGTGGTAATCTCCGAGCA | C12 |
| HvCHS I265F FP<br>HvCHS I265F RP | C3 | GAGGCGGGCTTAACGTTTCACCTGCTGAAAG<br>CTTTCAGCAGGTGAAACGTTAAGCCCGCCTC | C13 |
| PhCHS T197A FP<br>PhCHS T197A RP | C1 | GATTACCGCGGTGGCCTTCCGTGGCCC<br>GGGCCACGGAAGGCCACCGCGGTAATC | C14 |
| HvCHS A228S FP<br>HvCHS A228S RP | C3 | GCAATCATCGGAtCCGATCCCGACC<br>GAGTTCCAATTGTGAATACCCAAAGGCTCAAACGC | C15 |
| HvCHS D231I FP<br>HvCHS D231I RP | C3 | CGGAGCCGATCCCATCCAATTAGACGAG<br>GAGTTCCAATTGTGAATACCCAAAGGCTCAAACGC | C16 |
| HvCHS Q232P FP<br>HvCHS Q232P RP | C3 | GCCGATCCCGACCCGTTAGACGAG<br>GAGTTCCAATTGTGAATACCCAAAGGCTCAAACGC | C17 |
| HvCHS L233G FP<br>HvCHS L233G RP | C3 | GATCCCGACCAAGGTGACGAGCAGC<br>GAGTTCCAATTGTGAATACCCAAAGGCTCAAACGC | C18 |
| HvCHS D234V FP<br>HvCHS D234V RP | C3 | CCCGACCAATTAGTTGAGCAGCCGG<br>GAGTTCCAATTGTGAATACCCAAAGGCTCAAACGC | C19 |
| HvCHS Q232P_D234V_FP<br>HvCHS Q232P_D234V_RP | C15 | CCCGACCAATTAGTTGAGCAGCCGG<br><br>GAGTTCCAATTGTGAATACCCAAAGGCTCAAACGC | C20 |

*5’ [PHO] means 5' phosphorylation of primers

**Table S2.** Apparent Michaelis-Menten kinetic parameters for turnover of feruloyl-CoA by HvCHS wildtype and HvCHS (Q232P, D234V) at a fixed concentration of malonyl-CoA of 300 µM.

| **Enzyme** | **HvCHS** | **HvCHS (Q232P, D234V)** |
|---|---|---|
| **Best-fit values** | | |
| Et, µM | 0.05 | 0.05 |
| $k_{cat}$, $min^{-1}$ | 2.40 | 2.46 |
| $K_m$, µM | 4.18 | 2.63 |
| $V_{max}$, µM/min | 0.12 | 0.12 |
| **95% CI (profile likelihood)** | | |
| $k_{cat}$, min-1 | 2.16 to 2.72 | 2.24 to 2.72 |
| $K_m$, µM | 3,06 to 5,60 | 1,99 to 3,40 |
| **Goodness of Fit** | | |
| Degrees of Freedom | 16 | 16 |
| R squared | 0,9195 | 0,9054 |
| Sum of Squares | 0,001094 | 0,0009365 |
| Sy.x | 0,008271 | 0,007651 |

**Table S3.** Structure alignment of the new HvCHS structure (8B32) with CHS structures in the PDB with the Dali server.

| rank | PDB- chain | Z score | rmsd | lali | nres | %id | enzyme name with ligand | donor species |
|---|---|---|---|---|---|---|---|---|
| 1 | 4yjy-A | 68.2 | 0.5 | 387 | 393 | 79 | Chalcone Synthase 1 | *Oryza sativa* |
| 2 | 4wum-C | 68.2 | 0.4 | 386 | 389 | 73 | Chalcone Synthase | *Freesia hybrida* |
| 7 | 1cgk-A | 67.7 | 0.5 | 386 | 387 | 72 | Chalcone Synthase 2 with naringenin | *Medicago sativa* |
| 8 | 7bur-A | 67.7 | 0.5 | 385 | 388 | 74 | Chalcone Synthase 1 | *Gkycine max* (L.) |
| 9 | 1bi5-A | 67.6 | 0.5 | 386 | 389 | 72 | Chalcone Synthase 2 | *Medicago sativa* |
| 10 | 1i86-A | 67.6 | 0.5 | 386 | 389 | 72 | Chalcone Synthase 2 G256A | *Medicago sativa* |
| 11 | 1cgz-A | 67.6 | 0.5 | 386 | 387 | 72 | Chalcone Synthase 2 with resveratrol | *Medicago sativa* |
| 12 | 1d6f-A | 67.6 | 0.5 | 386 | 389 | 72 | Chalcone Synthase 2 C164A | *Medicago sativa* |
| 13 | 1bq6-A | 67.6 | 0.5 | 386 | 388 | 72 | Chalcone Synthase 2 with CoA | *Medicago sativa* |
| 14 | 1cml-A | 67.5 | 0.6 | 386 | 389 | 72 | Chalcone Synthase 2 with malonyl-CoA | *Medicago sativa* |

### Supporting Figures

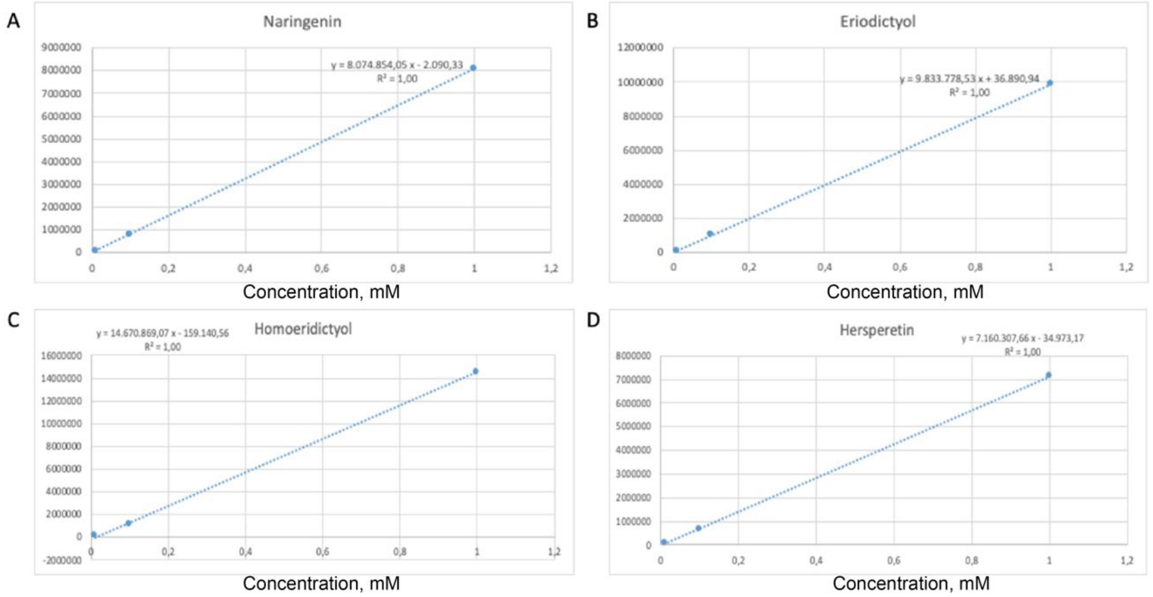

**Figure S1.** Calibration plot of flavonoids (A: naringenin, B: eriodictyol, C: homoeriodictyol, and D: hesperetin) dissolved in DMSO and analyzed by HPLC. The compounds were detected at 288 nm and the analysis was performed as described in the method section. The range of calibration curve was 0.01 mM to 1 mM.

A

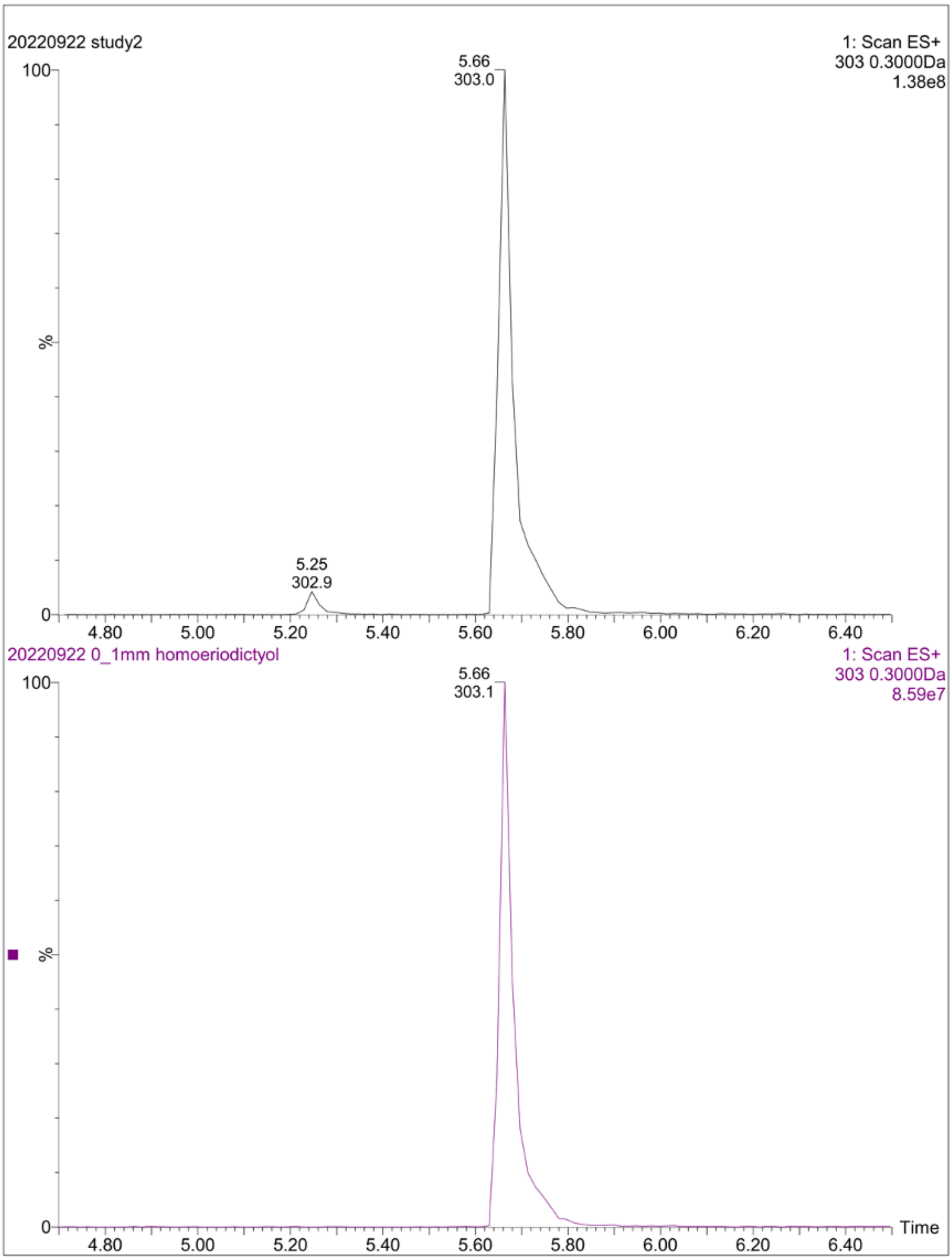

20220922 study2
1: Scan ES+
303 0.3000Da
1.38e8
5.66
303.0
5.25
302.9
100
%
0
4.80
5.00
5.20
5.40
5.60
5.80
6.00
6.20
6.40
20220922 0_1mm homoeriodictyol
1: Scan ES+
303 0.3000Da
8.59e7
5.66
303.1
Time

B

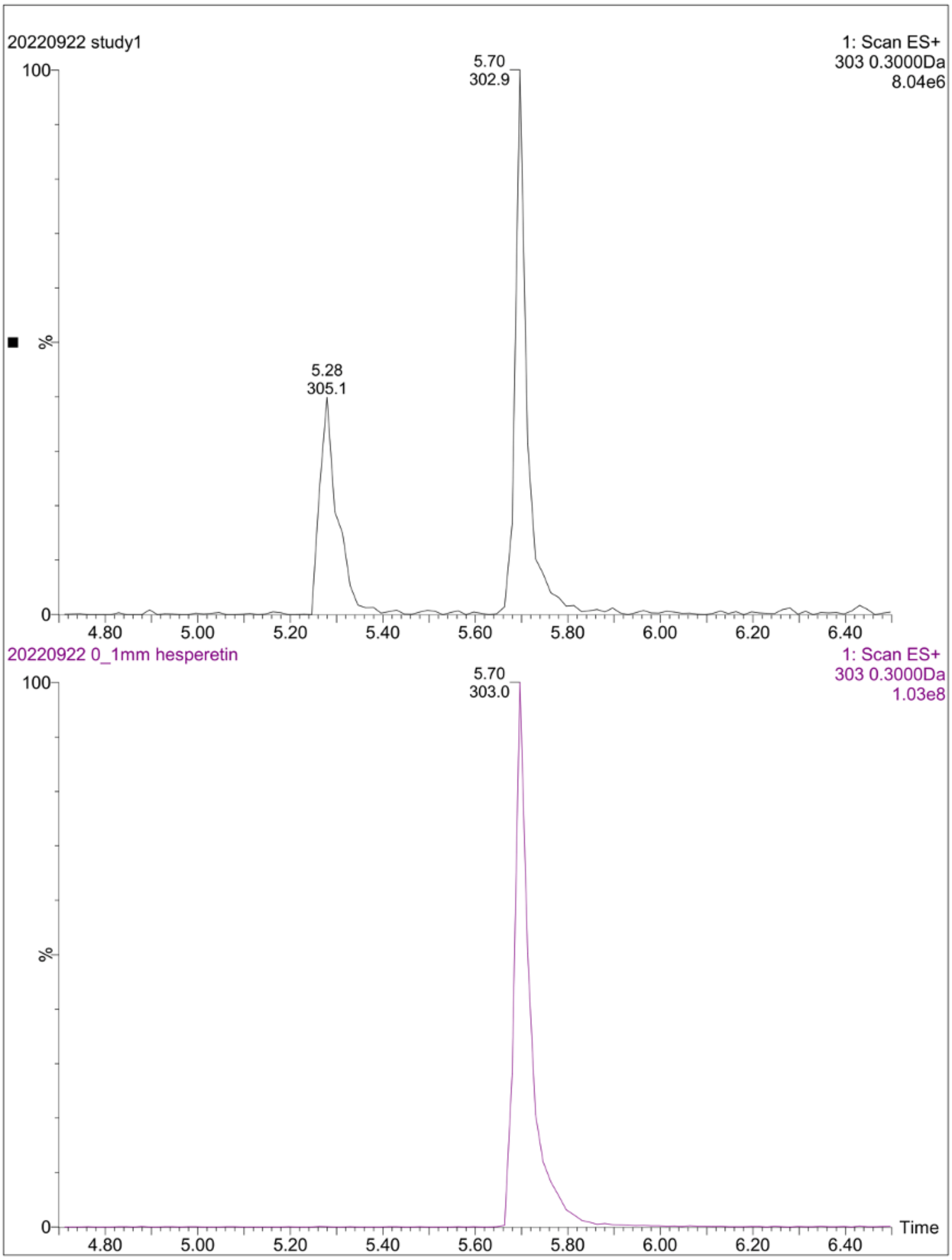

**Figure S2.** Extracted ion chromatograms at *m/z 303* [M+H⁺] in positive ion channel of the fermentation products with strain s2 expressing HvCHS wild type (black) and the analytical standards (pink). (A) homoeriodictyol (B) hesperetin.

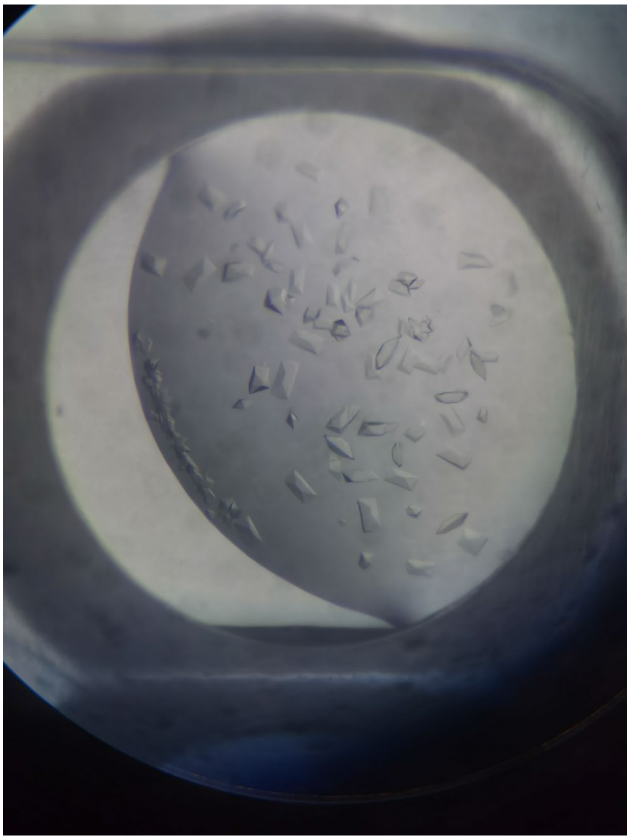

**Figure S3.** Micrograph of HvCHS crystals in the crystallization experiment. Multiple orthorhombic crystals grew in a clear, sitting drop containing 1µl protein (10mg/ml stock concentration) and 1µl reservoir solution (0.1 M MES/Imidazole pH6.5; 0.03 M $MgCl_2$, 0.03 M $CaCl_2$; 16% (v/v) glycerol, 8% (v/v) PEG4000) in 1-2 days at 4°C.

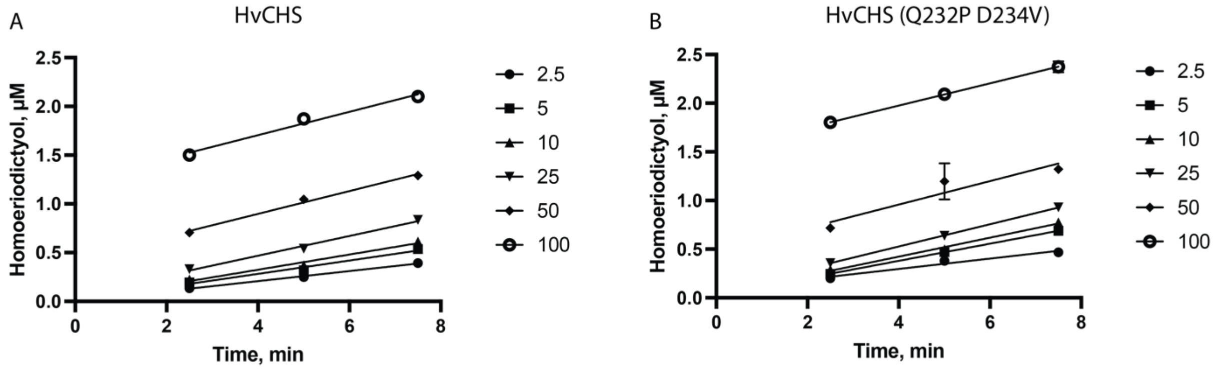

**Figure S4.** Time progress curves underlying the steady-state kinetics analysis. Concentration of homoeriodictyol obtained in *in vitro* turnovers catalyzed by A) HvCHS wild type and B) HvCHS double mutant in the presence of varying concentrations of feruloyl-CoA (2.5, 5, 10, 25, 50, and 100 µM) and a fixed concentration of malonyl-CoA (300 µM). Data points represent mean +/- SD, n=3, line represents linear regression to determine the apparent initial velocities for each substrate concentration.

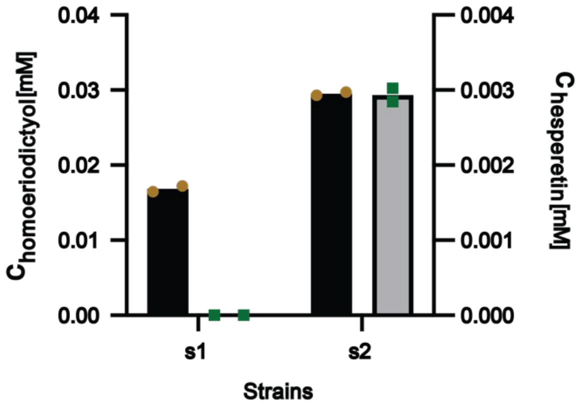

**Figure S5.** Comparison of different CHS and 4CL variants (s1: PhCHS from *Petunia hybrida* and 4CL from *Petroselinum crispum* and s2: CHS from *Hordeum vulgare* and 4CL from *Oryza sativa*). Small scale fermentation for s1 and s2. 1 mM ferulic acid, and isoferulic acid were added as precursors. Samples were taken after 32 h fermentation and analyzed by HPLC-MS. Each experiment was duplicated.

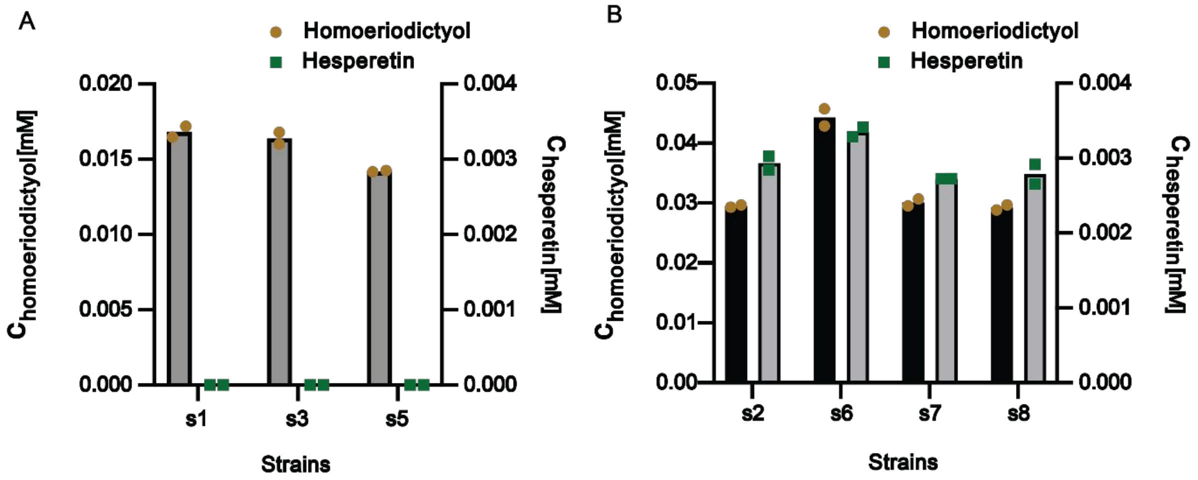

**Figure S6.** Comparison of different 4CL variants: A) Pc4CL variants (s1 Pc4CL wildtype, s3 Pc4CL del V342 variant, and s5 Pc4CL Q214A variant) and B) Os4CL variants (s2 Os4CL wildtype, s6 Os4CL del V340 variant, s7 Os4CL Q212A variant, and s8 Os4CL S242A variant). Small scale fermentation for those variants. 1 mM ferulic acid, and isoferulic acid were added as precursor. Samples were taken after 32 h fermentation and analyzed by HPLC-MS. Each experiment was duplicated.

```
                                                                                |
HvCHS      -MAAVRLKEVRMAQRAEGLATVLAIGTAVPANCVYQATYPDYYFRVTKSEHLADLKEKFQ   59
OsCHS      MAAAVTVEEVRRAQRAEGPATVLAIGTATPANCVYQADYPDYYFRITKSEHMVELKEKFK   60
FhCHS      ---MVNVEEIRKAQRAEGPAAILAIGTATPPNAIEQSEYPDYYFRVTNSEDKVELKEKFK   57
PhCHS      ---MVTVEEYRKAQRAEGPATVMAIGTATPSNCVDQSTYPDFYFRITNSEHKTDLKEKFK   57
GmCHS      ---MVSVEEIRKAQRAEGPATVMAIGTATPPNCVDQSTYPDYYFRITNSEHMTELKEKFK   57
MsCHS      ---MVSVSEIRKAQRAEGPATILAIGTANPANCVEQSTYPDFYFKITNSEHKTELKEKFQ   57
               * :.* * ****** *::***** * *.: *: ***:**::*:**. .:*****:

           ||  |
HvCHS      RMCDKSMIRKRHMHLTEEILIKNPKICAHMETSLDARHAIALVEVPKLGQGAAEKAIKEW   119
OsCHS      RMCDKSQIRKRYMHLTEEILQENPNMCAYMAPSLDARQDIVVVEVPKLGKAAAQKAIKEW   120
FhCHS      RMCEKSMIKKRYLYLTEDILKENPNVCAYMATSLDARQDMVVVEVPKLGKEAATRAIKEW   117
PhCHS      RMCEKSMIKKRYMHLTEEILKENPSMCEYMAPSLDARQDIVVVEVPKLGKEAAQKAIKEW   117
GmCHS      RMCDKSMIKKRYMYLNEEILKENPSVCAYMAPSLDARQDMVVVEVPKLGKEAATKAIKEW   117
MsCHS      RMCDKSMIKRRYMYLTEEILKENPNVCEYMAPSLDARQDMVVVEVPRLGKEAAVKAIKEW   117
           ***:** *::*:::*.*:** :**.:* :*  *****: :.:****:**: ** :*****

                          |
HvCHS      GQPLSKITHLVFCTTSGVDMPGADYQLTKLLGLSPTVKRLMMYQQGCFGGATVLRLAKDI   179
OsCHS      GQPRSRITHLVFCTTSGVDMPGADYQLAKMLGLRPNVSRLMMYQQGCFAGGTVLRVAKDL   180
FhCHS      GQPKSKITHLVFCTTSGVDMPGADYQLTKLLGLRPSVKRLMMYQQGCFAGGTVLRLAKDL   177
PhCHS      GQPKSKITHLVFCTTSGVDMPGCDYQLTKLLGLRPSVKRLMMYQQGCFAGGTVLRLAKDL   177
GmCHS      GQPKSKITHLIFCTTSGVDMPGADYQLTKLLGLRPSVKRYMMYQQGCFAGGTVLRLAKDL   177
MsCHS      GQPKSKITHLIVCTTSGVDMPGADYQLTKLLGLRPYVKRYMMYQQGCFAGGTVLRLAKDL   177
           *** *:****:.**********.****:*:*** * *.* ********.*.****:***:

                         ||| |            |     |||           * *
HvCHS      AENNRGARVLVVCSEITAMAFRGPCKSHLDSLVGHALFGDGAAAAIIGADPDQLDEQPVF   239
OsCHS      AENNRGARVLAVCSEITAVTFRGPSESHLDSMVGQALFGDGAAAVIVGSDPDEAVERPLF   240
FhCHS      AENNRGARVLVVCSEITAVTFRGPSESHLDSLVGQALFGDGAAALIVGSDAIEGIERPIF   237
PhCHS      AENNKGARVLVVCSEITAVTFRGPNDTHLDSLVGQALFGDGAGAIIIGSDPIPGVERPLF   237
GmCHS      AENNKGARVLVVCSEITAVTFRGPTDTHLDSLVGQALFGDGAAAVIVGSDPLP-VEKPLF   236
MsCHS      AENNKGARVLVVCSEVTAVTFRGPSDTHLDSLVGQALFGDGAAALIVGSDPVPEIEKPIF   237
           ****:*****.****:**::****  .:****:**:*******.* *:*:*     *:*:*

                         |||       ||||  |    |
HvCHS      QLVSASQTILPESEGAIDGHLTEAGLTIHLLKDVPGLISENIEQALEDAFEPLGIHNWNS   299
OsCHS      QMVSASQTILPDSEGAIDGHLREVGLTFHLLKDVPGLISKNIERALGDAFTPLGISDWNS   300
FhCHS      EMVSAAQTILPDSEGAIDGHLREVGLTFHLLKDVPGIISKNIEKSLEEAFKPLGITDYNS   297
PhCHS      ELVSAAQTLLPDSHGAIDGHLREVGLTFHLLKDVPGLISKNIEKSLEEAFKPLGISDWNS   297
GmCHS      QLVWTAQTILPDSEGAIDGHLREVGLTFHLLKDVPGLISKNIEKALVEAFQPLGISDYNS   296
MsCHS      EMVWTAQTIAPDSEGAIDGHLREAGLTFHLLKDVPGIVSKNITKALVEAFEPLGISDYNS   297
           ::* ::**: *:*.******* *.***:********::*:** ::* :** **** ::**

              ||||                          |
HvCHS      IFWIAHPGGPAILDRVEDRVGLDKKRMRASREVLSEYGNMSSASVLFVLDVMRKSSAKDG   359
OsCHS      IFWVAHPGGPAILDQVEAKVGLDKERMRATRHVLSEYGNMSSACVLFILDEMRKRSAEDG   360
FhCHS      LFWIAHPGGPAILDQVEAKIGLKPEKLRATRHVLSEYGNMSSACVLFILEEMRKKSAEEK   357
PhCHS      LFWIAHPGGPAILDQVEIKLGLKPEKLKATRNVLSNYGNMSSACVLFILDEMRKASAKEG   357
GmCHS      IFWIAHPGGPAILDQVEAKLGLKPEKMEATRHVLSEYGNMSSACVLFILDQMRKKSIENG   356
MsCHS      IFWIAHPGGPAILDQVEQKLALKPEKMNATREVLSEYGNMSSACVLFILDEMRKKSTQNG   357
           :**:**********:** ::.*. :::.*:*.***:*******.***:*: *** * ::

                             |
HvCHS      LATTGEGKDWGVLFGFGPGLTVETLVLHSVPVPVPTAASA   399
OsCHS      HATTGEGMDWGVLFGFGPGLTVETVVLHSVPITAGAAA--   398
FhCHS      NGTTGEGLEWGVLFGFGPGLTVETVVLHSVEA--------   389
PhCHS      LGTTGEGLEWGVLFGFGPGLTVETVVLHSVAT--------   389
GmCHS      LGTTGEGLDWGVLFGFGPGLTVETVVLRSVTL--------   388
MsCHS      LKTTGEGLEWGVLFGFGPGLTIETVVLRSVAI--------   389
             ***** :************:**:**:**
```

**Figure S7.** Multiple sequence alignments of those proteins that are structurally most similar to HvCHS as identified by Dali search against the PDB. Gray box: catalytic cysteine; pink bars: flavonoid binding residues, black bars: malonyl-CoA binding residues, purple box: active site residues that were mutated in this study; blue box: surface loop residues that were mutated in this study with stars marking the most beneficial mutations; color coding within black box according to CLUSTAL W color coding.

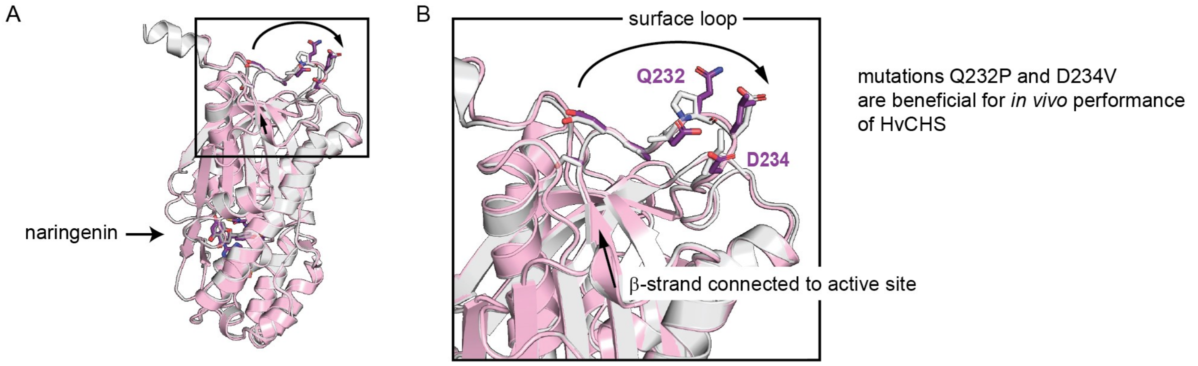

**Figure S8.** Illustration of surface loop in HvCHS (PDB: 8B35, pink) compared to MsCHS (PDB: 1CGK, grey), which appears to modulate HvCHS expression level and affinity to ferulic acid. The Q232P and D234V single and double point mutations appear to be beneficial for the *in vitro* and *in vivo* performance of the enzyme. Atom coloring in stick representation: oxygen=red, nitrogen=blue, phosphorous=orange, carbon=color matches the cartoon color of the corresponding structure. Compared to Fig. 2A, the model is rotated by 180° along the y-axis.
